# The prevalence of copy number increase at multiallelic CNVs associated with cave colonization in Mexican tetra (*Astyanax mexicanus*)

**DOI:** 10.1101/2023.11.10.566513

**Authors:** Ivan Pokrovac, Nicolas Rohner, Željka Pezer

**Affiliations:** Ruđer Bošković Institute, Zagreb, Croatia; Stowers Institute for Medical Research, Kansas City, MO, USA

**Keywords:** copy number variation, cave colonization, parallel evolution, ecological adaptation, gene duplication

## Abstract

Copy number variation is a common contributor to phenotypic diversity, yet its involvement in ecological adaptation is not easily discerned. Instances of parallelly evolving populations of the same species in a similar environment marked by strong selective pressures present opportunities to study the role of copy number variants (CNVs) in adaptation. By identifying CNVs that repeatedly occur in multiple populations of the derived ecotype and are not (or are rarely) present in the populations of the ancestral ecotype, the association of such CNVs with adaptation to the novel environment can be inferred. We used this paradigm to identify CNVs associated with recurrent adaptation of the Mexican tetra (*Astyanax mexicanus*) to cave environment. Using a read-depth approach, we detected CNVs from previously re-sequenced genomes of 44 individuals belonging to two ancestral surface and three derived cave populations. We identified 102 genes and 292 genomic regions that repeatedly diverge in copy number between the two ecotypes and occupy 0.8% of the reference genome. Functional analysis revealed their association with processes previously recognized to be relevant for adaptation, such as vision, immunity, oxygen consumption, metabolism, and neural function and we propose that these variants have been selected for in the cave or surface waters. The majority of the ecotype-divergent CNVs are multiallelic and display copy-number increases in cave fish compared to surface fish. Our findings suggest that multiallelic CNVs - including gene duplications, and divergence in copy number provide a fast route to produce novel phenotypes associated with adaptation to subterranean life.

**Significance Statement:** Duplications and deletions of genomic sequences occur frequently within a population. Such inter-individual difference in the amount of genetic material is known as copy number variation and is associated with differences in phenotypic traits. Despite its pervasiveness, the evolutionary impact of copy number variation in complex organisms is difficult to discern, primarily due to the infeasibility of setting up evolutionary experiments in the laboratory. Instances of multiple populations that evolved the same traits in similar environments represent naturally occurring evolutionary experiments and valuable opportunities to study the molecular basis of adaptation. Mexican tetra represents such a system and we use the genomes of surface and cave-dwelling populations to study the role of copy number variation in the recurrent cave adaptation.

## Introduction

Biological evolution operates on phenotypic and genetic variation. Structural variation is a dominant form of genetic variation in terms of genome proportion, population frequency, and taxonomic ubiquity. It refers to the inter-individual variation in the orientation, position, or copy number of a genomic sequence. The latter covers deletions and amplifications of sequences, collectively termed copy number variants (CNVs). CNVs affect both coding and noncoding regions and thus may alter phenotype in various ways. For example, by affecting complete genes CNVs may cause changes in gene dosage (Maron et al. 2013; Handsaker et al. 2015). They can modulate the expression of individual genes by affecting enhancers or promoters, or reorganize whole regulatory networks by striking at a single critical transcription factor (Vickrey et al. 2018; Yuste-Lisbona et al. 2020). CNVs in exons cause changes in gene structure, consequently changing the structure and efficiency of protein products (Boettger et al. 2016). With such prevalence and prolific influence on phenotype, CNVs are considered to have a large impact on phenotypic diversity.

Despite their pervasiveness among genomes and taxa, it is not clear to what extent CNVs contribute to adaptation. Even in the most extensively studied species - humans, no consensus has been reached yet and the assumptions range from neutral evolutionary processes acting on the majority of CNVs to the significant contribution of adaptive evolution (Iskow et al. 2012; Saitou et al. 2022). Multiple factors complicate the reconciliation between studies such as the heterogeneity of CNVs in terms of type, size, genomic context, and mutation rate, as well as technical difficulties and limitations pertaining to the choice of methodology (reviewed in Pokrovac and Pezer 2022). Moreover, experimental evolution as a tool is generally not feasible for more complex multicellular organisms. The dynamics of CNVs at macro- and microevolutionary scales are therefore inferred from comparative genomics and population-scale data, respectively, by examining evolutionary signatures, such as divergence patterns and CNV frequencies. In this respect, the instances of multiple populations with similar traits in similar environments represent precious opportunities for studying the evolution by natural selection, because they provide an element of reproducibility: if the trait repeatedly occurs in parallel, it is unlikely to have been driven by genetic drift, but rather it (and the underlying genetic variant) evolved multiple times as a response to a common selection pressure associated with habitat similarity (Rundle et al. 2000). CNVs are surprisingly understudied in the context of parallel evolution and recurrent adaptation. The only system that has received somewhat more attention in this regard is freshwater colonization by marine fish (Hirase et al. 2014; Lowe et al. 2018; Ishikawa et al. 2022). These studies identified CNVs, including changes in gene copy number (CN), that underly adaptation to freshwater environments, by analyzing data from multiple ancestral and derived populations.

We here use population-scale genomic data from Mexican tetra (*Astyanax mexicanus*) to investigate the contribution of CNVs to adaptation to cave environments. *A. mexicanus* is a fish species that exists in two forms: surface-dwelling, which inhabits lakes and rivers throughout Mexico and southern Texas, and cave-dwelling, which can be found in waters of multiple caves in northeastern Mexico (Gross 2012). The molecular data suggests that there are two lineages of *A. mexicanus* - new and old lineage, which separated 150,000 - 300,000 years ago (Herman et al. 2018). The cave form is considered derived from ancestral surface populations that invaded caves on more occasions 10,000 - 100,000 years ago and these transitions occurred in both lineages (Fumey et al. 2018). Despite some degree of gene flow from surface populations and evidence of reticulate evolution, troglomorphic traits are maintained in cave populations (Herman et al. 2018) indicating the presence of strong selection pressures to cope with environmental challenges such as constant darkness, reduced food availability, low oxygen level, and low parasite diversity. These are the key forces that drive the evolution of cave-derived traits, such as eye degeneration (Moran et al. 2015), loss of pigment (Bilandžija et al. 2013), sleep loss (Duboué et al. 2011; Jaggard et al. 2018), increase in appetite and starvation resistance (Aspiras et al. 2015), increased fat accumulation (Xiong et al. 2018), insulin resistance (Riddle et al. 2018), enlarged hypothalamus (Menuet et al. 2007), changes in behavior (Yoshizawa et al. 2010; Elipot et al. 2013; Kowalko et al. 2013), larger number and size of erythrocytes (Boggs et al. 2022; van der Weele and Jeffery 2022), and shift in the immune investment strategy (Peuß et al. 2020). The strong driving selective factors and distinctive derived traits, a well-known population history, and the existence of multiple ancestral and derived populations make *A. mexicanus* a convenient system for studying the role of CNVs and other structural variants in recurrent adaptation.

We use available genomic data from 44 fish belonging to three cave and two surface populations (Herman et al. 2018) to detect CNVs and discover signatures of natural selection associated with the transition to underground waters. To discover CNVs from NGS data, we employ CNVpytor (Suvakov et al. 2021), a Python extension of CNVnator and a tool that identifies deletions and duplications from regions with a lack or excess of mapped reads, respectively (Abyzov et al. 2011). Such a read-depth-based approach is highly accurate at estimating diploid CNs from short-read sequencing data, is robust to interindividual differences in genome coverage, and performs well in repetitive regions (Abyzov et al. 2011; Pezer et al. 2015; Kosugi et al. 2019; Garg et al. 2021). These features make it suitable for comparing genome-wide patterns of CN between populations. We use it to identify genes and genomic regions whose copy numbers diverge in parallel between cave and surface populations. Our findings highlight the role of multiallelic CNVs and the copy number increase in rapid adaptation to cave environment.

## Results

### Population diversity

By using CNVpytor we detected between 3,001 and 12,262 CNVs per animal (Table S2 in Supplementary Material). There was no significant difference between lineages or ecotypes in the total number of detected CNVs. New lineage animals contained a higher proportion of duplications than animals of the old lineage (Welch t-test, p-value 9×10^-12^, Cohen’s D 2.9): approximately 8% of all calls in Choy and Molino fish are duplications, whereas they constitute on average 4% of all calls in populations belonging to the old lineage. The higher duplication-to-deletion ratio in new-lineage animals can be explained by the fact that the AstMex3 reference is based on the fish from the Choy population, which belongs to the new lineage. Genomes are expected to share more of their content within the same lineage than between lineages, *i.e.* fewer regions in the reference genome are expected to be missing in the new-lineage genomes than in the genomes of old-lineage fish. There was no major difference in the proportion of duplications between surface and cave ecotypes (Welch t-test, p-value 0.36).

The extent of shared genetic variation between individuals can provide clues on genetic diversity. In order to estimate relative genetic diversity within and between populations, we analyzed the number of overlapping CNV calls between any two individuals in our dataset. Predicted CNV call breakpoints are defined by genomic coordinates in the reference genome and the distance between the breakpoints defines CNV length. If the genomic position of a CNV call in one genome overlaps with the position of a call in another genome, and they do so over a minimum of 50% of both lengths, these calls are defined as a shared CNV between the two genomes. Based on such a definition, we find that any two samples share on average 1,946 CNVs, corresponding to 34% of all CNVs in a single pairwise comparison. Individuals are the most similar to one another in the Molino population which belongs to the cave-dwelling new lineage, where they share 3,448 CNVs on average (65%). This finding suggests the lowest genetic diversity in Molino population, in accord with previous observations based on analyses of SNP data (Bradic et al. 2013; Herman et al. 2018). Río Choy population (surface-dwelling new lineage) is the most diverse, with individual pairs sharing on average 1,198 CNVs (36%). The pattern of shared CNVs clusters populations according to their lineages (Figure 1A) but groups the Rascon surface and Tinaja cave population together. This disagrees with the previously proposed phylogeny based on SNP data in which cave populations Tinaja and Pachón form a monophyletic sister group to Rascon surface population (Herman et al. 2018).

**Figure 1.**
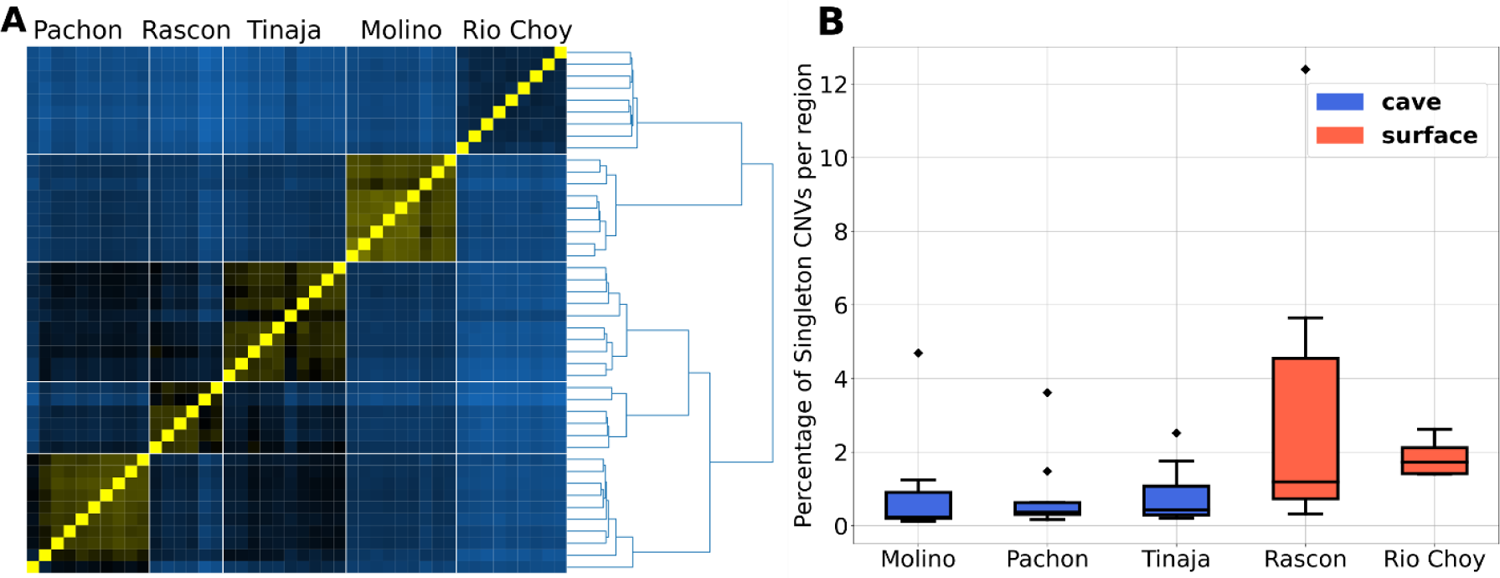
Genetic diversity analysis based on shared and singleton CNVs. **A)** Distance matrix based on the average number of shared CNVs between two genomes. Samples are shown in the same order from right to left as from top to bottom. Identity is depicted as the brightest yellow, whereas maximum dissimilarity (the smallest number of shared CNVs in the dataset) is shown by the lightest blue. Distance matrix was subjected to Ward’s method of hierarchical clustering. **B)** Distribution of singleton CNVs proportions by population.

In order to identify CNV loci that are specific to populations, we defined copy number variable regions (CNVRs) as regions enclosed by genomic coordinates of merged calls from all individuals of the same population. This enabled us to determine population-private CNV loci, *i.e.* calls present within CNVRs in at least one individual of a single population whereas absent from all other populations (Table 1). We find 4,257 cave-specific and 4,728 surface-specific CNVRs. The proportion of private CNVRs that are found in single animals is greater in surface fish (65%), than in cave fish (41%). This is also evident in comparisons of individual surface populations (72% in Río Choy and 68% in Rascon) with individual cave populations (45% - 62%). A single private CNVR contains on average five CNVs in cave fish and only three in surface animals (Table 1). These analyses suggest that cave fish more often share the same CNV, consistent with pairwise similarity analysis based on the number of shared CNVs (Figure 1A). It was proposed that most genetic variation in the caves results from standing genetic variation from the ancestral surface stock and possible gene flow between the populations (Bradic et al. 2012). Hence, genetic variation in cave animals is expected to largely represent a subset of surface variation. The finding of a similar number of cave-specific CNVRs and surface-specific CNVRs is therefore surprising. There was no significant difference between cave and surface in the functional content of ecotype-specific CNVRs (Figure S1). Majority of the genes that overlapped these CNVRs were associated with numerous signaling and metabolic pathways to similar extent in both ecotypes.

**Table 1.** Count of CNV regions that are private to population, ecotype and lineage.

| **Group** | **Private CNVs** | **Private CNVRs** | **CNVRs\* with single CNV** | **CNVs per CNVR\* (average)** |
| --- | --- | --- | --- | --- |
| Molino | 3,577 | 1,215 | 551 (45%) | 5 |
| Pachón | 1,770 | 807 | 502 (62%) | 4 |
| Tinaja | 3,142 | 1,266 | 623 (49%) | 4 |
| Río Choy | 1,264 | 830 | 596 (72%) | 3 |
| Rascon | 5,649 | 3,527 | 2,383 (68%) | 3 |
| cave | 15,016 | 4,257 | 1,750 (41%) | 5 |
| surface | 8,243 | 4,728 | 3,056 (65%) | 3 |
| new lineage | 6,346 | 2,326 | 1,197 (52%) | 5 |
| old lineage | 35,583 | 8,752 | 3,633 (42%) | 6 |

\* Numbers refer to CNVRs that are private to population, ecotype, or lineage, as indicated by the first column

We next analyzed the proportion of singleton CNVs, defined as CNVs detected in only one animal and having no overlap with CNVs in other samples of the whole dataset. We detected between 5 and 1,537 singletons per animal - corresponding to 0.12% - 12.53% of all CNVs. The proportion of singletons, relative to the number of all detected CNVs within an individual, is higher in surface populations than in cave populations (Figure 1B). This suggests that surface fish have larger diversity compared to cave fish and agrees with previous analyses based on microsatellite and SNP data (Bradic et al. 2012; Herman et al. 2018). The singleton proportion as well as the CNV presence-absence pattern suggest low genetic diversity within the Pachón population (Figure 1). This population is characterized by small size and stronger isolation, and our results align with the low polymorphism observed from microsatellite data (Legendre et al. 2023).

To estimate the fraction of the AstMex3 genome that is copy number variable in the whole set, we merged calls across all samples into CNVRs. We found in total 15,088 CNVRs on assembled chromosomes, comprising cumulatively 260.2 Mbp of sequence. Compared to the total sequence length of assembled chromosomes (1,321 Mbp), this translates into 19.7% of the reference sequence being copy number variable in natural *A. mexicanus* populations. We discover that 10,035 CNVRs overlap genes (including protein-coding genes, lncRNA, rRNA, tRNA, and all other annotated gene classes) and occupy 213 Mbp in total, whereas the rest 5,053 (occupying 48 Mbp in total) can be considered purely noncoding CNVRs. The larger proportion of CNVRs associated with genes may be explained by the high proportion of gene-encoding sequences present in the reference assembly: as much as 63% of the assembled sequence is annotated as genes in the AstMex3.

### Population-specific patterns of gene copy number

We detected in total 2,819 unique protein-coding genes that are entirely spanned by a CNV in at least one sample. We refer to this set of genes as CNV genes (Table S3 in Supplementary_Tables.xlsx). The average length of a CNV gene is 9,263 bp (median 5,335 bp), which is substantially shorter than the average length of all annotated protein-coding genes (34,441 bp; median 13,069 bp). Over a third of CNV genes (978) are uncharacterized, *i.e.* of yet-unknown function, which is a four-fold increase compared to the proportion of uncharacterized protein-coding genes in the whole assembly (8.5%). This enrichment of uncharacterized genes in the set of CNV genes could be a consequence of their generally short length (7.3 kbp on average in the whole genome), such that smaller genes are more likely to entirely reside within CNVs than longer genes. In order to test this, we performed permutation analyses. Coordinates of CNV calls were shuffled randomly, keeping the distribution of CNV lengths as is in the true data. In each permutation, the average length of CNV genes and the proportion of uncharacterized CNV genes were calculated. Based on 100 such permutations, we would expect to find about 10% of uncharacterized genes within the set of CNV genes, which is very close to the proportion seen in the whole AstMex3. Moreover, we would expect their average length to be around 15.7 kbp, which is lower than the genome average (34.4 kbp) yet substantially higher than the true CNV genes set (9.3 kbp). CNV genes are therefore 3.5 times more likely to be uncharacterized and 1.7 times shorter than expected. A previous study on three-spined stickleback fish demonstrated that CNVs are enriched for evolutionary new genes which are generally shorter and for which no evidence of homology in other species has been found (Chain et al. 2014). Their function is often unknown; hence these genes are described as “uncharacterized”. Notably, given the recency of the AstMex3 assembly and the associated genome annotations, some of the genes labeled as “uncharacterized” may not actually be true orphans or lineage-specific genes, but may instead represent genes that have orthologs in other species, of yet undetermined function. However, the lack of such characterization, short gene length, and the strong enrichment compared to expectations based on permutations in our study, provide further support to the hypothesis that CNVs are enriched for young genes (Chain et al. 2014).

By using the *-genotype* option in CNVpytor we determined the copy number of every CNV gene in each sample (Table S3). Based on gene copy numbers, the samples cluster by their geographic location (Figure 2A and 2B). Interestingly, the two surface populations, new-lineage Río Choy and old-lineage Rascon, are closer than expected, given that the split between the two lineages happened at least 200,000 years ago (Herman et al. 2018). Moreover, hierarchical clustering suggests that Rascon surface and Tinaja cave fish are the most similar based on the pattern of gene copy number and that the two populations form a sister clade with the Río Choy surface population (Figure 2B).

**Figure 2.**
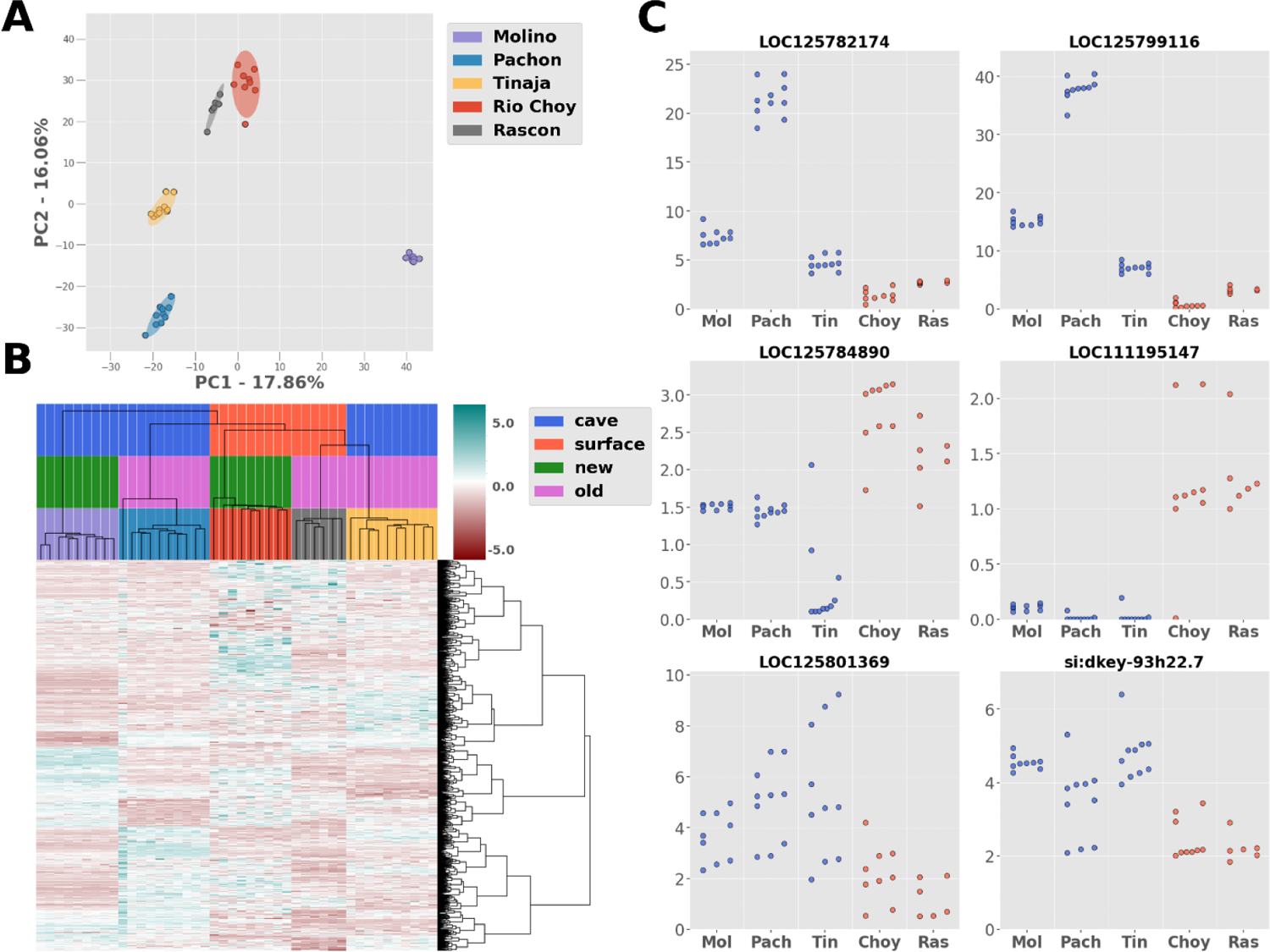
Population specific pattern of gene copy number variation. **A)** Plot of the first two principal components of PCA based on copy numbers of 2,819 CNV genes. Confidence (95%) ellipses are shown around sample clusters. **B)** Heatmap of normalized copy numbers of the 2,819 CNV genes with hierarchical clustering based on genes (rows) and samples (columns) using Ward’s method. Values are normalized by row (gene) such that the mean of every row is 0 and its standard deviation is 1. The heatmap visually represents relative differences in copy number of a particular gene between samples, ranging from lowest (red) to highest (green) values. The samples are colored as in A) by population, and by ecotype and lineage as indicated in the legend on the right. **C)** Copy numbers of several genes with significant differences between cave (blue) and surface (red) individuals. Gene symbol is indicated on top of each plot. Mol - Molino; Pach - Pachón; Tin - Tinaja; Choy - Río Choy; Ras - Rascon.

To find genes with significant differences in CN between ecotypes, we performed appropriate statistical tests for all combinations of cave-surface population pairs, as well as for comparison of all cave versus all surface individuals (see Methods section for details). Based on this approach, we found that 102 out of 2,819 CNV genes were significantly different in copy number between cave and surface ecotypes (Table S3). Over half (65) are predicted genes of unknown function among which 89% (58 genes) have an average copy number higher in cave than in surface fish. For example, we find between 4 and 24 copies of the *LOC125782174* in cave individuals, whereas the same gene exists in 1-3 copies in the surface fish (Figure 2C). Similarly, we find up to 4 copies of *LOC125799116* in genomes sampled in surface waters and 6-40 copies in cave genomes. Among the genes that are characterized in this set, some have either lower or higher average copy number in cave fish genomes compared to surface, and play a part in different processes such as (innate) immunity (*LOC111188594* - fucolectin-1-like; *LOC111194616* - polymeric immunoglobulin receptor-like; *LOC111195410* - E3 ubiquitin-protein ligase DTX3L; *LOC125785782*, *LOC125785783*, *LOC111188451* and *LOC125785616* - B-cell receptor CD22-like; *LOC111197671* - C-type lectin domain family 4 member E-like; *LOC125784871*, *LOC125785663* and *LOC125784856* - NLR family CARD domain-containing protein 3-like), oxygen transport (*LOC111191628*, *LOC111196759*, *LOC111191630* and *LOC103027764* - encoding hemoglobin subunits; *LOC125787068* - scavenger receptor cysteine-rich type 1 protein M130-like), and lipid metabolism (*LOC103026892* - 60 kDa lysophospholipase; *LOC125799429* - phospholipase B-like 1; *LOC111195147* - apolipoprotein L3-like). For example, gene *LOC125784890* is predicted to encode trace amine-associated receptor 13c on chromosome 20. This gene belongs to a family of vertebrate olfactory receptor genes and its copy number is reduced in cave genomes compared to surface fish (Figure 2C). Similarly, gene *LOC111195147* predicted to encode apolipoprotein L3-like seems to be completely deleted in cave genomes and is present mainly in one or two copies in surface populations (Figure 2C).

Apolipoprotein L3 in humans is implicated in the movement of lipids (including cholesterol) within cytoplasm, and the binding of lipids to organelles (Gene database - National Library of Medicine; Gene ID: 80833). Gene *LOC125801369* which encodes a protein similar to zinc finger protein 501 on chromosome 4 is present in 3-9 copies in most cave fish samples, whereas it is present in 1-2 copies in the majority of surface genomes. Similarly, the copy number of gene *si:dkey-93h22.7* is amplified in cave fish populations (Figure 2C). This gene encodes golgin subfamily A member 6-like protein 22 and its expression is restricted to testis in humans (Gene database - National Library of Medicine; Gene ID: 440243).

Similarly, we found 157 genes with significant differences in their CN between lineages (Table S3). Among the 74 uncharacterized genes in this set, 40 had an average copy number higher in new lineage compared to old lineage animals. Several annotated genes stand out as being specifically amplified or deleted in only one lineage. For example, we find a substantially higher copy number of the *ERVFC1* gene annotated on chromosome 16 in old-lineage animals. This gene encodes endogenous retroviral envelope protein and is present in 1-3 copies in Choy and Molino fish, and in 6-27 copies in genomes of Rascon, Tinaja, and Pachón fish (Figure S2). Instances of gene *PGBD4* annotated on chromosomes 1, 2, 10, 19, and 20 are present mainly as one copy per diploid in old-lineage animals and as two or three copies in the new-lineage animals. This gene belongs to the family of piggyBac transposable element-derived (*PGBD*) genes that are found in diverse animals (Sarkar et al. 2003). In humans, its expression is enhanced in skeletal muscle, spermatids, and immune cells (Human Protein Atlas; Uhlén et al. 2015). Similarly, homologs of gene *SMAD3* annotated at different genomic locations on chromosomes 4, 5, 11, 15, 17 and 22, are reduced to a single copy per diploid on average in fish of the old lineage, compared to on average three copies in the new lineage. In humans, this gene encodes a widely expressed transcription factor with roles in many cellular processes such as cell proliferation, cell movement, and apoptosis (Human Protein Atlas; Uhlén et al. 2015).

### Ecotype-divergent CNVs at genes

Differences in copy number between populations can result in phenotypic differences between them. If genes affected by divergent CNVs are associated with particular biological processes, this may indicate that these processes have also diverged between the populations. To identify such processes that might be specifically targeted for copy number divergence between surface and cave fish, we explored the functional context of genes that are frequently affected by CNVs in single ecotypes.

We find a total of 9,187 protein-coding genes that are overlapped by CNVs. At 1,653 (18%) of the genes, we detected CNVs in only one sample, and at 476 (5%) genes CNVs are found in all 44 analyzed genomes (Figure S3). Of the 9,187 genes, 15% (1,407) were affected by CNVs exclusively in cave (Molino, Pachón, Tinaja) populations (Table S4) and 19% (1,726) in surface (Rascon, Río Choy) populations (Table S5). Within this set of ecotype-specific events, the largest proportion of the genes were affected in only one animal (42% or 589 genes in the cave, and 62% or 1,064 genes in the surface). A much smaller fraction overlapped CNVs in multiple animals and in at least one animal per population, *i.e.* 4% (58 genes) in the cave and 9% (163 genes) in the surface fish. Genes that are the most frequently affected by ecotype-specific events are genes that are associated with immune response. For example, genes such as *LOC125785663* (NLR family CARD domain-containing protein 3-like), *LOC111196508* (scavenger receptor cysteine-rich type 1 protein M130), *LOC103031898* (deleted in malignant brain tumors 1 protein), and *pikfyve* (phosphoinositide kinase, FYVE finger containing) are associated with innate immunity, inflammatory response and antiviral defense. CNVs in these genes are detected in 13-28 cave individuals (of the 29 in total; Table S4). Similarly, genes implicated in these processes are found to overlap the most frequent surface-specific events: *LOC103031476* (NACHT, LRR and PYD domains-containing protein 3), *rnf41* (ring finger protein 41), *LOC125801137* (E3 SUMO-protein ligase ZBED1-like) and *LOC125782699* (scavenger receptor cysteine-rich type 1 protein M130-like) are affected by CNVs in 9-13 out of 15 surface individuals (Table S5). Within the set of cave-specific events with the highest frequency (>30%) in analyzed populations, we find genes associated with processes such as visual perception (*rgrb* - retinal G protein-coupled receptor b; *LOC111194948* - TOG array regulator of axonemal microtubules protein 1; *map2* - microtubule-associated protein 2; *opn8a* - opsin 8 group member a; *LOC107197208* - complement C1q-like protein 3), genes encoding hemoglobin subunits (*LOC111191630* - hemoglobin subunit beta-2-like; *LOC111191631* - hemoglobin embryonic subunit alpha; *LOC111191628* - hemoglobin embryonic subunit alpha), and genes implicated in neurological functions (*plppr3a* - phospholipid phosphatase related 3a; *LOC103030484* - ras-related protein Rab-26; *rab3c* - RAS oncogene family member). Genes associated with these processes are either not or are not frequently affected by CNVs in surface populations (Tables S4 and S5).

Ecotype-divergent CNVs can affect genes along their whole length, as described for CNV genes above. For example, gene *LOC125785663* appears to be either entirely deleted or reduced to a single copy in cave individuals (Figure 3, Table S3). An ecotype-divergent CNV gene can be affected by both deletions and duplications in different individuals, such as the gene *LOC111194948*, which appears to be either deleted or amplified in the cave genomes, while present mainly in two copies in surface individuals (Figure 3, Table S3). CNVs can affect only a part of a gene at high frequency, such as in the case of *rgrb* and *rnf41*. In all analyzed cave genomes, a region from exon 4 to 5 of *rgrb* is deleted. Similarly, a region encompassing the first intron and the first two exons of *rnf41* is amplified in all analyzed surface fish (Figure 3).

**Figure 3.**
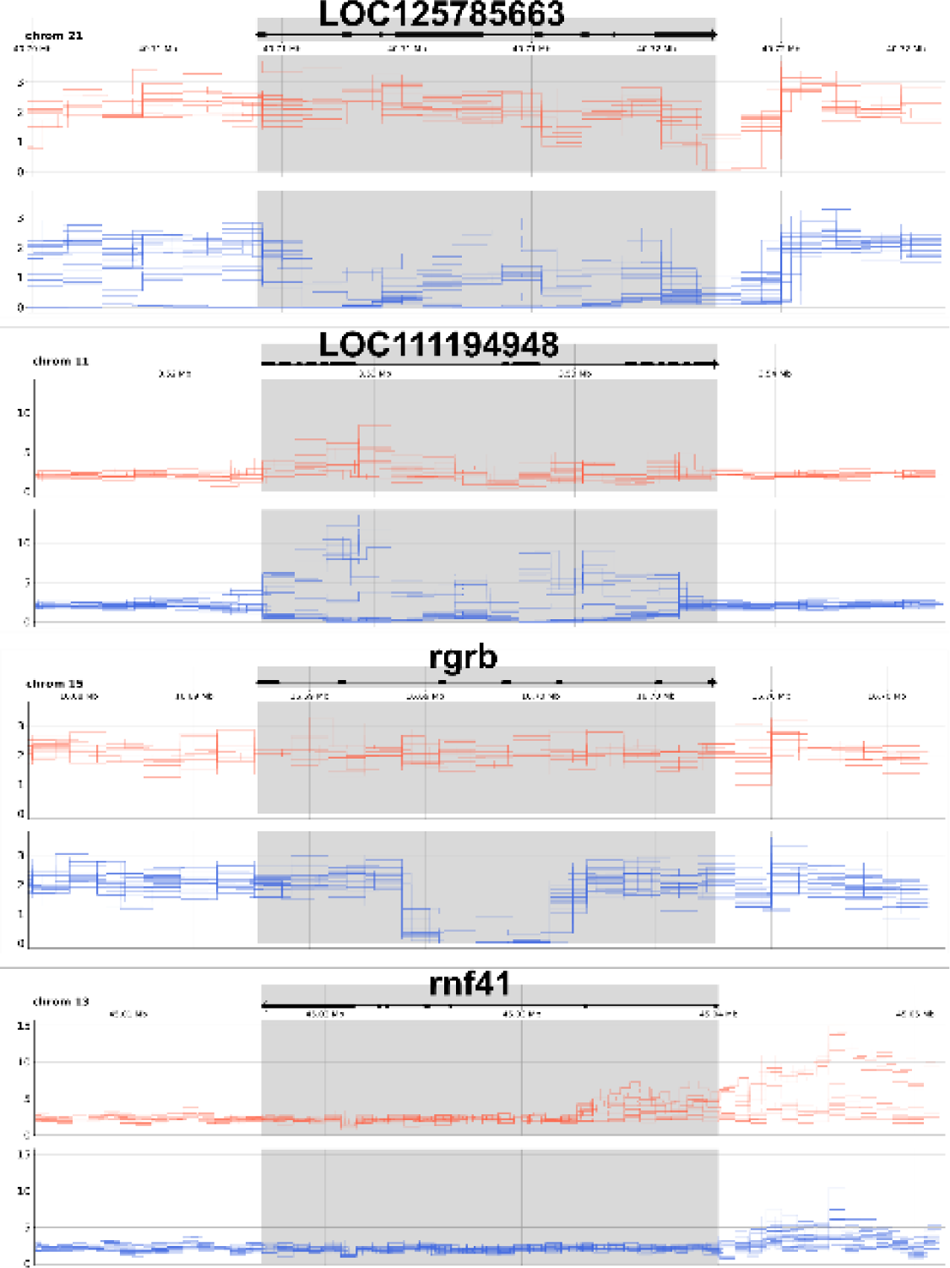
Read depth in the regions of several ecotype-divergent CNVs at genes. Read depth is shown per bin for cave (blue) and surface (orange) fish, as calculated by CNVpytor and normalized to represent the copy number per diploid. Gene structure and orientation are shown by arrows on top of which gene symbols are indicated. Gene position is highlighted in gray.

### Ecotype-divergent CNVRs

Of the 15,088 CNVRs detected in the whole set, 292 showed significant differences in copy number between cave and surface animals (Wilcoxon rank sum test, adjusted pval < 0.01), in all combinations of surface-cave comparisons as well as in comparison of all surface fish with all cave fish (Table S6). These constitute in total 10.47 Mbp, equaling 0.8% of the reference genome. Almost a third of these CNVRs (87) are noncoding, whereas the majority (205) overlap or contain one or more genes (Table S6). Average copy numbers of these genomic regions range much wider in cave fish than in surface fish. Of the CNVRs that show significant differences in average CN between ecotypes, the largest majority varies only slightly in surface fish, existing mainly in one or two copies per diploid. The same genomic regions in cave fish genomes are amplified up to 34 copies on average (Figure 4A). Two-thirds of these regions (195/292) exist at higher copy numbers in cave fish compared to surface, whereas only 97 CNVRs are estimated to have lower copy numbers in cave fish.

**Figure 4.**
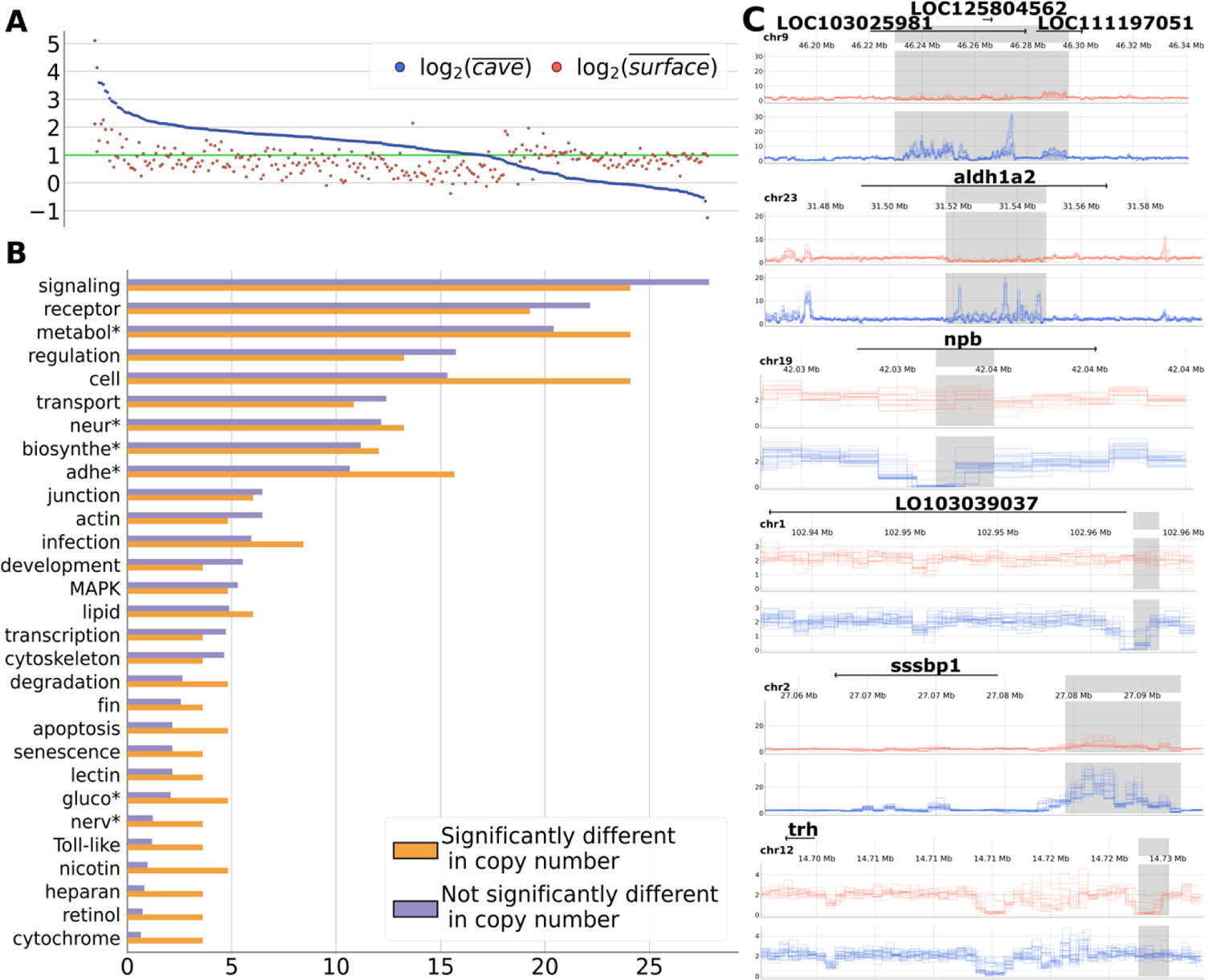
Genomic regions divergent in copy number between surface and cave fish. **A)** Genomic regions that are significantly different in their copy number between the two ecotypes. All 292 CNVRs are shown on the X axis, where each position represents one CNVR. Every CNVR is genotyped with CNVpytor and the average CN per individual is shown as blue and orange dots for cave and surface fish, respectively. CNVRs are ordered by average copy number in cave fish from highest on the left to the lowest on the right. Copy numbers are shown as log2-transformed values on y-axis; the green horizontal line depicts value of two copies per diploid. **B)** The most frequently occurring terms associated with genes that overlap CNVRs, analyzed separately for regions that significantly differ (orange) or do not differ (blue) in copy number between cave and surface fish. Terms are derived from biological process and pathway annotations in DAVID tool. Complete list of terms associated with words marked by asterisks is given in Table S8. Proportions are shown as percentages of the number of genes associated with a term relative to the total number of genes that have assigned annotations in DAVID. **C)** Read depth around several CNVRs (highlighted in gray) with divergent copy numbers between ecotypes. Gene positions and orientations are shown by arrows on top of which gene symbols are indicated. Read depth is plotted per bin for cave (blue) and surface (red) fish, as calculated by CNVpytor and normalized to represent the copy number per diploid.

We analyzed functional annotations of genes overlapping CNVRs with significant differences in their copy number between ecotypes. Of the 460 genes in the 205 regions, only 83 had associated annotations for biological processes or pathways (Table S7). About half of these genes were associated with numerous signaling and metabolic pathways, including MAPK, Toll-like and C-type lectin signaling pathways (Figure 4B and Table S8). Other highly represented categories included transport of substances, regulation of different processes, and processes related to nervous system functioning such as neuroactive ligand-receptor interaction and neurotransmitter transport. These categories were also present at similarly high proportions in genomic regions that did not significantly differ in copy number between ecotypes (Figure 4B).

However, some processes seem to be more represented in genomic regions with divergent copy number between cave and surface animals. They include categories associated with cell adhesion, apoptosis, glucose metabolism, as well as metabolisms of cytochrome, retinol and nicotinate/nicotinamide, and the degradation of branched-chain amino acids (Figure 4B and Table S8). Notably, among divergent CNVRs, we find many that intersect genes potentially involved in biological processes that have been shown to change upon adaptation to darkness, such as vision, development, and behavior. For example, genes *LOC111197051* and *LOC103025981* are annotated in the AstMex3 assembly as two copies of the same gene that encodes guanylyl cyclase-activating protein 2, which is involved in phototransduction. These genes overlap a ∼66 kb long genomic region that is affected by duplications only in cave fish (Figure 4C). Similarly, a 31 kb genomic region is amplified in cave fish that overlaps gene *aldh1a2* (Figure 4C). This gene encodes retinaldehyde dehydrogenase 2, which is an enzyme essential for proper embryonal morphogenesis (Niederreither et al. 2002). A small, ∼1 kb region within gene *npb* appears to exist in cave fish as a single copy per diploid (Figure 4C). This gene encodes neuropeptide B, which is expressed in the central nervous system and implicated in regulating feeding behavior (Singh and Davenport 2006).

Noncoding genomic regions that are significantly different in copy number between ecotypes may be relevant for adaptation if they contain regulatory elements. Such alterations may cause differences in the regulation of specific processes associated with nearby genes. To explore this idea, we extracted the closest gene to each of the 87 divergent noncoding CNVRs and analyzed their functions. Although only 27 genes had associated annotations for biological processes or pathways, they mainly agree with the analysis of gene-encoding CNVRs described above (Table S9). Many of these genes can be associated with processes that are known to change upon adaptation to the cave environment. For example, a gene that encodes cytochrome c oxidase subunit 4 (*LOC103039037*) is located ∼350 bp downstream of a ∼1.4 kb region that is deleted in cave fish, whereas the same region exists as two copies per diploid in surface fish (Figure 4C). This gene is associated with oxidative phosphorylation and cardiac muscle contraction, and the cave-specific deletion of the CNVR next to this gene may reflect differences in its regulation as an adaptation to hypoxic conditions in caves. Another alteration that might be associated with adaptation to reduced oxygen levels in a subterranean environment is the amplification of a ∼8 kb genomic segment, about 5 kb upstream of *ssbp1*. This gene is important for mitochondrial biogenesis and it is tempting to speculate that 9-17 copies of the CNVR near this gene in cave fish might be responsible for the altered regulation of mitochondria production as a possible route to compensate for hypoxia (Gutsaeva et al. 2008; Gamboa and Andrad 2009). Interestingly, a small region (∼2 kb) located ∼27 kb upstream of *trh* gene is found at lower copy number in surface fish compared to cave fish (Figure 4C). This gene is expressed in hypothalamic neurons as thyrotropin-releasing hormone that has major roles in many biological processes including metabolic activity, thermoregulation, locomotor activity, pain perception, and sleep regulation (Wozniak and Quinnell 2015).

In summary, functional analysis of the genes in or near CNVRs suggests that the same biological processes may diverge through copy number variation in genes as well as in noncoding regions, potentially affecting regulatory elements. Products of genes associated with such CNVRs function in processes that are considered relevant for adaptation to the underground environment such as metabolism, visual processing, morphogenesis, oxygen consumption, and behavior.

### Evolutionary implications

On average, 57% of all detected CNVs overlap protein-coding genes (Figure 5A), 12% of which affect whole genes. The second most prominent group of genes that are affected are genes that encode long noncoding RNAs, with approximately 10% of all detected events overlapping them. About a third of all CNVs have no overlap with any of the categories related to gene features (Figure S4).

**Figure 5.**
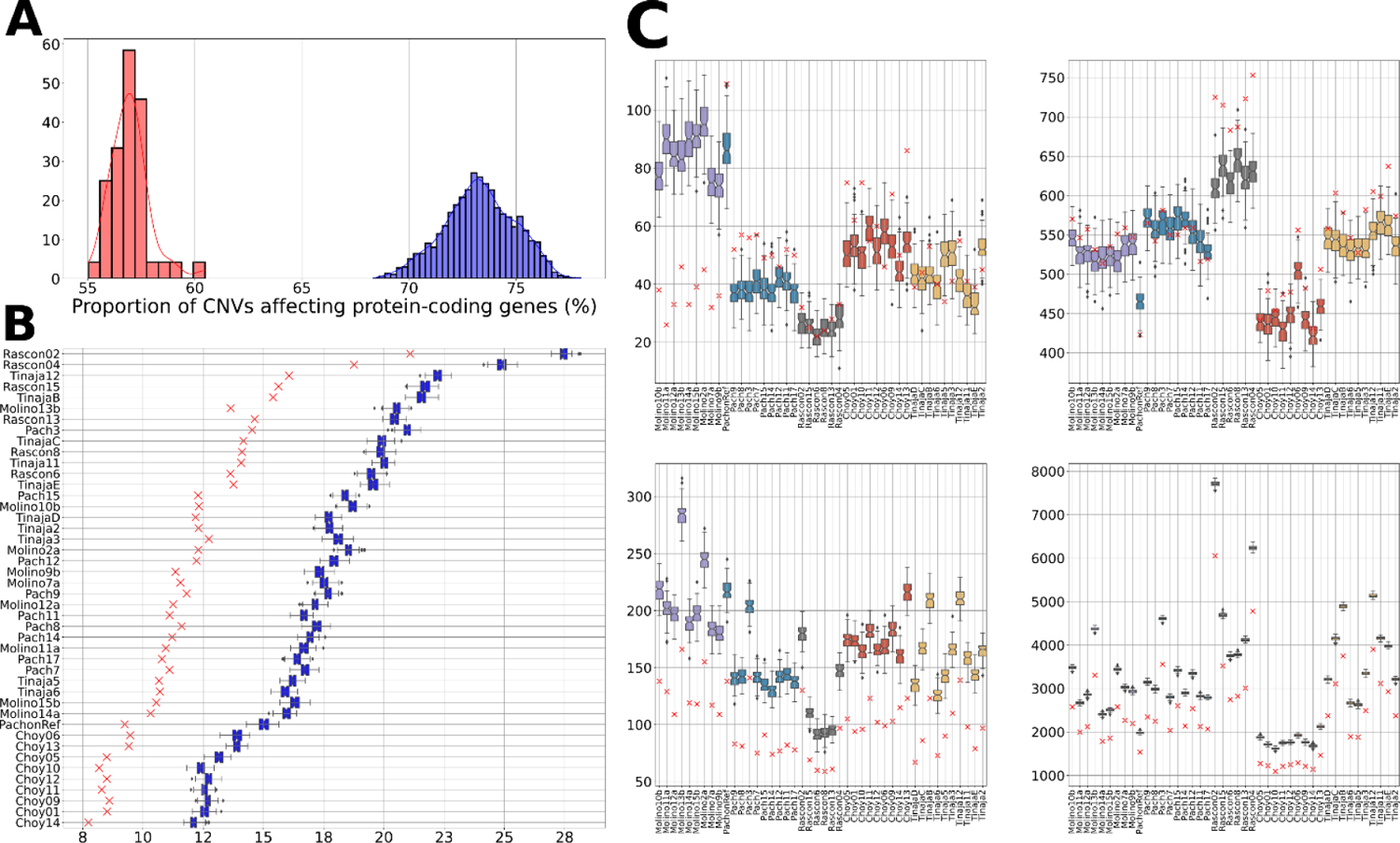
**A)** Distribution of percentage of CNVs affecting protein-coding genes across all 44 analyzed individuals, for the true (red) and permuted (blue) data. **B)** Proportion of protein-coding genes affected by CNV calls in true data (red crosses) and the distribution of expected proportions based on permuted data (blue boxplots). Data is shown by individuals, indicated by sample names on the left. **C)** Number of duplications (left plots) and deletions (right plots) that affect protein-coding genes entirely (upper plots) or partially (lower plots). Values are indicated per individual, as red crosses for true data and as boxplots for permuted data, representing the distributions of values based on 100 permutations. Boxplots are colored by population.

Permutations of calls in analyzed samples can provide clues to the mode of evolution that acts on CNVs affecting particular gene categories. For example, if the detected proportion of CNVs affecting a particular group of genes is similar to the expected proportion, those CNVs can be considered to represent neutral variation. In contrast, if the expected proportion is higher or lower than the detected proportion, CNVs affecting such category of genes may be under purifying or positive selection, respectively. In order to see if the detected proportions of CNVs overlapping different gene features are expected by chance, or if there is some bias for or against CNVs, we permuted the calls and analyzed their overlap with each of the annotated gene categories (Figure 5, Figure S5). Analysis of permutated data suggests that between 69% and 78% of all CNVs would affect protein-coding genes by random chance (Figure 5A). Comparison with the detected proportions that range from 55% to 61% suggests that CNVs are generally biased away from protein-coding genes in all samples. This is further corroborated when the number of affected genes is considered: on average 18% of all protein-coding genes are expected to be affected by CNVs based on permutations, compared to the detected 12% of genes in the true data (Figure 5B).

The bias against CNVs in protein-coding genes is contributed by events affecting parts of genes, whereas complete genes appear to be duplicated or deleted in a neutral fashion (Figure 5C). For the most part, this is in agreement with previous findings in humans that suggested purifying selection acts against all types of structural variants that affect protein-coding genes, except complete duplications (Collins et al. 2020). Similarly, it was demonstrated that duplication events, rather than deletions, are more likely to include entire protein-coding genes in the stickleback fish genomes (Lowe et al. 2018). The apparently neutral variation of gene copy number in our dataset could explain the stratification of samples based on CNV genes (Figure 2), which roughly reflects the demographic history of these populations. Interestingly, we find two exceptions: Molino population, in which fewer gene duplications are detected than expected - suggesting purifying selection, and Rascon population, in which gene deletions seem to be subject to positive selection (Figure 5C, upper panels). To investigate if the observed bias against CNVs affecting part of genes is a function of the unusually large intron sizes in the Mexican tetra (Jakt et al. 2022), we analyzed the overlap of CNVs with exons and introns. The comparison of permuted and true data suggests that CNVs affecting a part of intron or exon are subject to neutral evolution (Figure 6). Interestingly, complete intron or exon duplications seem to be better tolerated than deletions. Therefore, the general bias against CNVs at protein-coding genes is contributed predominantly by selective constraints against deletions of complete exons or introns, whereas neutral evolutionary forces seem to shape variation in gene copy number.

**Figure 6.**
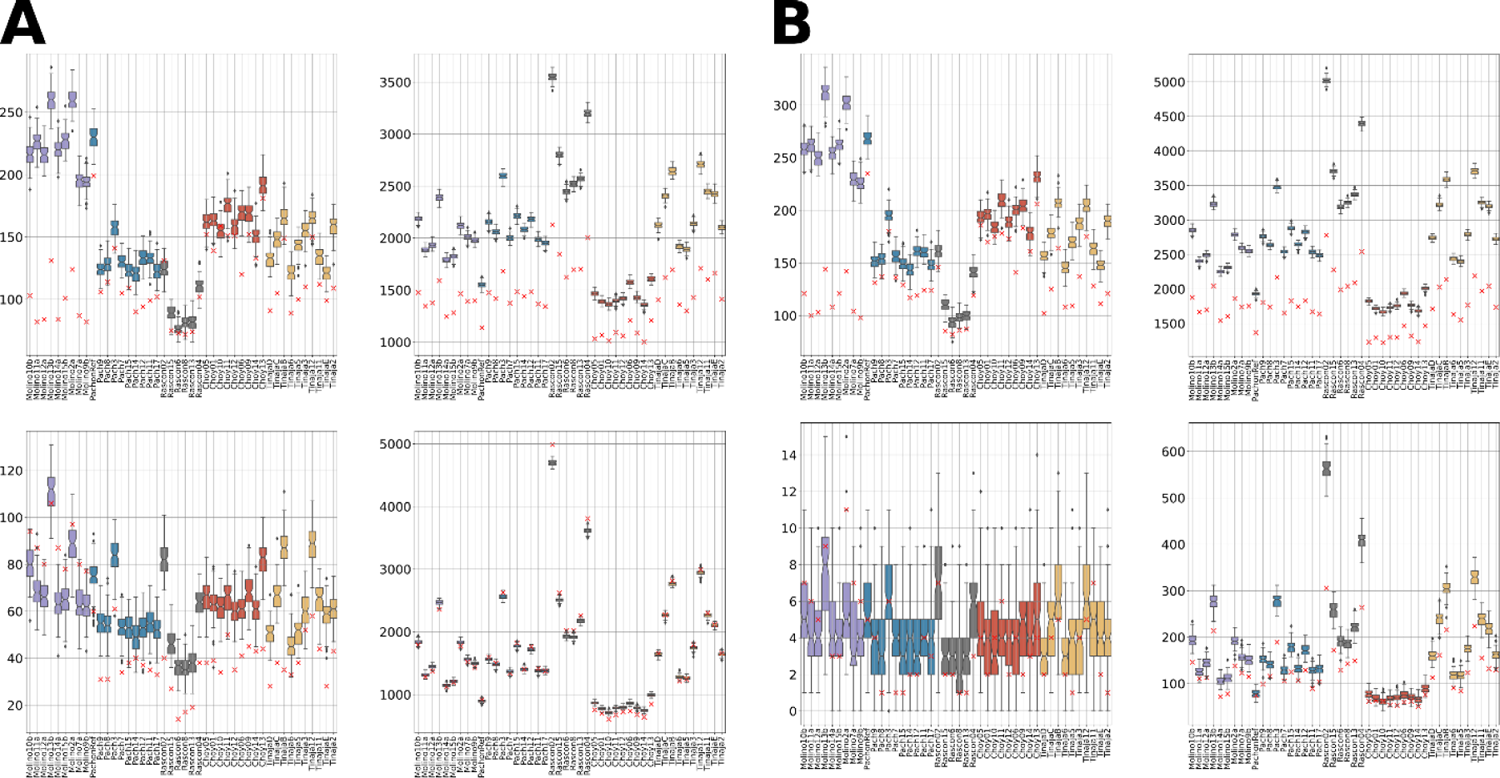
Number of duplications (left plots) and deletions (right plots) that affect. **A)** introns and **B)** exons entirely (upper plots) or partially (lower plots). Values are indicated per individual, as red crosses for true data and as boxplots for permuted data. Each boxplot represents a distribution of values based on 100 permutations. Boxplots are colored by population.

Based on comparisons of detected CNVs with permuted data (Figures 5, 6, and S5), we compiled the inferred mode of mutation of CNVs intersecting all major annotated gene categories in the AstMex3 assembly (Table 2). Interestingly, duplications of whole genes, regardless of gene category, seem to generally evolve neutrally, whereas deletions of complete genes are either neutral or under positive selection, such as in genes encoding lncRNAs, rRNAs, tRNAs, and pseudogenes. Surprisingly, only partial duplications and partial deletions of protein-coding genes seem to be subject to purifying selection (Figure 5B).

**Table 2.** Selection on duplications and deletions in gene categories inferred from comparisons of true and permuted data.

| Gene category | Category count<sup>2</sup> | Deletion complete<sup>1</sup> | Duplication complete<sup>1</sup> | Duplication partial<sup>1</sup> | Deletion partial<sup>1</sup> |
| --- | --- | --- | --- | --- | --- |
| protein-coding | 26,735 | 0* | 0* | - | - |
| pseudogene | 1,376 | + | 0* | 0 | + |
| lncRNA | 3,603 | + | 0* | 0 | 0 |
| snoRNA | 297 | 0 | 0 | N/A | N/A |
| snRNA | 1,314 | 0 | 0 | N/A | N/A |
| rRNA | 9,252 | + | 0 | N/A | N/A |
| tRNA | 9,987 | + | 0 | N/A | N/A |

<sup>1</sup> neutral variation: "0"; variation under negative selection: "-"; variation under positive selection: "+"; not applicable: "N/A" (partial overlap cannot be determined due to considerably smaller gene length compared to bin size used for CNV detection)
<sup>2</sup> number of annotated instances in the AstMex3 genome assembly
\* categories with exceptions to the indicated inference (for details, see Figure S5)

### Validity considerations

All CNVs in the dataset were detected and genotyped by CNVpytor, an extension of CNVnator developed in Python (Suvakov et al. 2021; Abyzov et al. 2011). CNVpytor has additional features and improved performance in terms of speed but retains CNVnator’s high sensitivity, low false-discovery rate, and high genotyping accuracy (Suvakov et al. 2021). The genotyping accuracy of its algorithm reaches 96% and is robust to differences in coverage (Abyzov et al. 2011; Pezer et al. 2015; Kosugi et al. 2019), rendering it one of the most commonly used tools for CNV detection based on the read-depth approach. Moreover, a recent study validated the genotyping performance of CNVnator in tandemly repetitive regions and found a high correlation (R^2^ = 0.81) between CNs deduced from short reads and long reads (Garg et al. 2021), demonstrating that CNVnator’s algorithm is reliable even in such problematic regions in which the mapping position of short reads cannot be determined with certainty. Our major findings are based on copy numbers genotyped by the CNVpytor. We rely on the reports of high accuracy described above as well as on the previously conducted validations in other studies that reported high correlation and concordance with copy numbers determined by different experimental and bioinformatic approaches (tan Nguyen et al. 2013; Yi et al. 2014; Shebanits et al. 2019; Meng et al. 2022).

We considered the possibility that our major findings could be influenced by the high proportion of mapped reads with low quality. For example, non-uniquely mappable reads could pile up at one particular position of a genotyped region and consequently inflate the inferred copy number of the whole region. We therefore assessed the proportion of reads with zero mapping quality (MAPQ=0) at each of the 292 ecotype-divergent CNVRs and the 102 ecotype-divergent CNV genes (Figure S6). At the largest majority of these genotyped loci, there were no low-quality reads mapped, and the proportion is smaller than 5% at approximately 90% of analyzed the loci. These numbers suggest that the effect of non-uniquely mapping reads on the genotyped copy numbers of CNVRs and CNV genes is insignificant.

## Discussion

Over the course of evolution, taxonomically diverse animals have successfully transitioned from surface to subterranean environments and thus converged to a set of adaptive traits such as loss of vision and pigment, and decline in metabolic rate. These species thus provide a natural setting for studying the molecular basis of adaptation to constant darkness and temperature, food scarcity, and low oxygen levels. Copy number variation as a form of genetic variation has not been sufficiently studied in this context, especially given that it has been recognized as a major contributor to phenotypic variation and rapid adaptation to novel and extreme environments (Kondrashov et al. 2002; Kondrashov 2012; Lye and Purugganan 2019; Rinker et al. 2019). A recent study addressed the role of CNVs in adaptation to subterranean environment at a macroevolutionary scale (Balart-García et al. 2023), yet a comprehensive exploration at the level of population is missing. We here considered the role of CNVs in the adaptation of the Mexican tetra to cave. By exploiting a set of genomic data previously generated from specimens collected at five distinct localities, we infer events and biological processes that may be under selection. The same genomes were previously studied by Warren et al. (2021), but only deletions were considered, and the study did not perform systematic analysis to detect divergence between ecotypes. Therefore, our study here represents the first comprehensive analysis of duplications and deletions from a micro-evolutionary perspective that identifies divergent CNVs associated with parallel cave colonization.

Considering all calls in analyzed individuals, we estimate that one-fifth of the reference sequence is subject to variation in copy number in natural *A. mexicanus* populations, the majority of which is contributed by CNVs affecting genes. However, it is important to note that only a third of the assembled AstMex3 consists of sequences outside genes. Previous studies in humans demonstrated that repetitive regions are particularly rich in structural variants, including CNVs (Huddleston et al., 2017; Audano et al., 2019; Ebert et al., 2021). By approaching the completion of the *A. mexicanus* reference genome assembly, especially with respect to repetitive sequences, we expect to see an increase in the proportion of CNVs in noncoding regions.

We observe strong stratification of CNVs between populations, in line with numerous studies in various species (Sudmant et al. 2015; Pezer et al. 2015; Xu et al. 2016; Dorant et al. 2020; Zhu et al. 2020; Jang et al. 2021; Solé et al. 2019; Yang et al. 2023). The profile of CNVs based on presence-absence patterns follows the previously established phylogenetic relationship between new and old lineage *A. mexicanus* populations based on SNPs (Herman et al. 2018).

However, within the old lineage, cave population Pachón is less similar to the Tinaja cave population, and the latter shares a larger proportion of CNVs with the Rascon surface population. This departure from SNP-based phylogenetic inference is even more obvious when CNV genes are considered: the hierarchical clustering based on gene copy number positioned the old lineage Pachón closer to the new lineage Molino population. These differences can be explained in the light of the elevated mutation rate of CNVs compared to SNPs (Zhang et al. 2009). Additionally, other factors may contribute to the observed patterns, such as the population size and the degree of migration between populations. Recent findings suggest that the Pachón population consists of only a few hundred individuals inhabiting a relatively isolated area, the direct consequences of which are low genetic polymorphism and limited gene flow from surface or other cave populations (Legendre et al. 2023). Accordingly, among the studied populations in our analyses, Pachón shows low diversity at the level of CNVs. Therefore, factors such as small population size and isolation of the Pachón population combined with the high CNV mutation rate may have increased the effect of genetic drift on CNVs genome-wide, resulting in an unexpected position of Pachón on the phylogenetic tree.

We used multiple approaches to estimate the relative CNV diversity of analyzed populations, and the results consistently indicate lower genetic diversity of cave populations compared to surface. Such finding agrees with previous studies based on SNPs and microsatellite data and can be explained by a combination of small effective population size, limited nutrient and space availability in caves, as well as possible bottleneck events (Bradic et al. 2012; Bradic et al. 2013; Herman et al. 2018). Most of the genetic variation in the caves is proposed to represent a subset of standing genetic variation from the ancestral surface stock (Bradic et al. 2012). However, we detected a high and comparable number of genomic regions that show copy number variation in an ecotype-specific manner, *i.e.* thousands of CNVs are detected in either cave or surface genomes but not both. Such observation raises a possibility that a great deal of copy number variation arises independently in both ecotypes. Nevertheless, these results are heavily dependent on the sample size, and many more genomes per ecotype would need to be screened for more reliable numbers and a firm interpretation of this finding.

To infer events that are likely under selection, we searched for genomic loci that satisfy all of the following criteria: 1) the region is divergent in copy number between surface and cave populations, 2) the divergence is significant in all cave-surface population-pair comparisons, and 3) the copy number change proceeds in the same direction in all population-pair comparisons. Such stringent criteria allowed us to identify almost three hundred genomic regions containing CNVs that may have been selected for in the cave or surface waters. These regions cumulatively account for nearly 1% of the genome. This proportion is well in line with the estimate obtained by a study that analyzed populations of three-spined sticklebacks in the context of freshwater colonization (Lowe et al. 2018). Within this set, we identified approximately one hundred genes with ecotype-divergent CNs, most of which show properties of young genes and are mainly amplified in cave populations. Similarly, two-thirds of all detected ecotype-divergent genomic regions have elevated copy numbers in cave (derived) genomes. Interestingly, gene copy number increase, rather than decrease, was also found to be dominant in the derived freshwater populations of sticklebacks (Hirase et al. 2014; Lowe et al. 2018; Ishikawa et al. 2022). These studies together with our findings suggest that larger gene-copy numbers may generally confer higher adaptive potential upon colonization of novel and extreme environments. An earlier analysis of human CNVs suggested that duplications are under less stringent evolutionary constraints than deletions, thus representing a larger target for adaptive selection (Sudmant et al. 2015). Moreover, duplications are more likely to show higher mutation rates due to the susceptibility to nonallelic homologous recombination between directly oriented duplicated sequences. This enables them to frequently change their copy-number state over a short time (Sudmant et al. 2015) and thus persist as multiallelic CNVs within a population. The majority of ecotype-divergent regions in our study are present at multiple copy-number states in the dataset and we show that many encompass complete genes. These multiallelic CNV genes may be particularly relevant in an evolving lineage and may increase fitness from the moment of their origin, most likely as a protein dosage effect in response to a changing environment (Kondrashov et al. 2002; Handsaker et al. 2015). Although we find a considerable proportion of cave-specific CNVs (detected only in the cave fish) in the whole dataset, the majority of the identified divergent regions show some copy number variation in the surface populations as well. Hence, the selection seems to draw from the pre-existing standing genetic variation in the ancestral populations, as the fastest route for adaptation to occur (Jones et al. 2012; Lai et al. 2019; Zong et al. 2021).

Many ecotype-divergent genomic regions in our study are associated with genes involved in biological processes that have previously been identified as important for the adaptation of *A. mexicanus* to life in caves. We find CNVs at or near various genes that are associated with the functioning of the central nervous system, visual processing, metabolism, oxygen consumption, and immune system, highlighting the involvement in environmental information processing as their common feature (Kondrashov et al. 2002). The parallel divergence of these variants in cave populations suggests their participation in physiological and behavioral responses to major challenges such as persistent darkness, low nutrient availability, low oxygen level, and differences in parasite composition.

## Conclusion

Our findings support the notion that gene duplications and divergence in copy number are key generators of evolutionary innovation associated with adaptation to subterranean life (Balart-García et al. 2023). In line with previous observations based on studies in other species, we suggest that CNVs contribute to phenotypic diversity and facilitate rapid ecological adaptation (Kondrashov et al. 2002; Sudmant et al. 2015; Rinker et al. 2019).

## Materials and Methods

### Data acquisition

The reference genome AstMex3_surface in FASTA format as well as annotation information and assembly report were downloaded from NCBI database, under RefSeq assembly accession GCF_023375975.1 (Warren et al. 2023). Illumina reads from *A. mexicanus* population resequencing data described in Herman et al. (2018) were downloaded from European Nucleotide Archive (SRA accession: PRJNA260715). Information about samples, including SRA run accession numbers is provided in Table S1. Sequencing data corresponding to the previous *A. mexicanus* genome assembly Astyanax_mexicanus-1.0.2 (GenBank accession GCA_004802775.1) was included as an additional sample of Pachón population (SRA accession: PRJNA533584). The data is based on 15 runs of Illumina HiSeq2000 on DNA isolated from heart, gill, and liver of a single female Pachón cave fish. Runs were joined and subsequently trimmed and cleaned using Trimmomatic (Bolger et al. 2014) and cutadapt (Martin, 2011) so that they pass quality control.

### Preprocessing and mapping

FASTQ files were quality checked with FastQC package (Andrews, 2010). Reads were mapped to the reference genome using Bowtie 2 (Langmead and Salzberg 2012) with default parameters. SAMtools (Li et al. 2009) was used to fix mate-pair information, sort the data, identify and mark duplicate reads, index the BAM files, and calculate the mean of per-base coverage.

### CNV calling

CNVs were detected with CNVpytor (Suvakov et al. 2021). The optimal bin size was set for each sample individually, such that the ratio of global RD mean to global RD standard deviation was between 4 and 5, corresponding to the relative standard deviation of global RD of 0.25 and 0.2, respectively. The bin size ranged from 500 to 800 bp. Copy numbers (CNs) were determined with CNVpytor by using the *-genotype* option.

### Downstream analyses

Principal component analysis and hierarchical agglomerative clustering were performed in Python programming language using *SciPy* and *sklearn* packages. Permutations were performed in Python *NumPy* package. Coordinates of CNV calls were shuffled randomly on the same chromosomes, while ensuring that CNV size distribution and duplications-to-deletions ratio matched those of the true data and that the annotated assembly gaps were avoided. CNV calls were intersected with specific features using *bioframe* package (Open2C et al. 2022).

Functional analysis of genes was performed by using the DAVID tool (Sherman et al. 2022). To assign biological processes and pathways, annotations from UP_KW_BIOLOGICAL_PROCESS, GOTERM_BP_DIRECT, and KEGG_PATHWAY were used. The most frequently occurring words from the DAVID output were extracted and compiled manually into a list of terms that encompass words or word roots. Such terms were used to analyze DAVID output by using the *grep* command in Linux. For example, the term “nerv*” was used to count all instances of the words *nerve, nervous,* and *innervation*. *Bedtools closest* was used to find the nearest genes to the noncoding CNVRs (Quinlan and Hall 2010).

To find genes with significant differences in CN between ecotypes or lineages, we combined the results of two approaches: 1) population-pair approach, in which Welch T-Tests were used in all combinations of a cave versus a surface population, or in all combinations of a new versus an old-lineage population; 2) bulk-comparison approach, in which Mann-Whitney test was used in comparison of all cave versus all surface individuals, or in comparison of all new-lineage versus all old-lineage individuals. The nonparametric Mann-Whitney test was chosen for bulk comparisons to account for multimodality in CN across the populations, given the population-specific profile of CNVs. We further adjusted p-values using the FDR Benjamin-Hochenberg method and FWER Holm’s method, setting the significance level at alpha 0.05. We considered the difference to be significant when the significance level criterion was met in both approaches. CNVPytor enables us to track two key metrics for genomic region analysis: total reads mapped, and unique reads mapped. To estimate the fraction of zero mapping quality reads (reads mapped to multiple regions), we calculate the difference between total and unique reads mapped to a region and divide it by the number of all reads mapped to the same region.

## Acknowledgments

This work was supported by the Croatian Science Foundation (grant UIP-2019-04-7898). NR is funded by the National Institutes of Health (NIH, grant R24OD030214). Data analysis was performed on the high-performance computing cluster at the University Computing Centre (SRCE), University of Zagreb.

## Author Contributions

ZP designed the research. NR contributed the data. IP and ZP performed the research and analyzed the data. ZP wrote the manuscript draft. All authors contributed to the final manuscript.

## References

Aspiras AC, Rohner N, Martineau B, Borowsky RL, Tabin CJ. Melanocortin 4 receptor mutations contribute to the adaptation of cavefish to nutrient-poor conditions. Proc Natl Acad Sci U S A. 2015 Aug 4;112(31):9668–73. doi: 10.1073/pnas.1510802112.

Abyzov A, Urban AE, Snyder M, Gerstein M. CNVnator: an approach to discover, genotype, and characterize typical and atypical CNVs from family and population genome sequencing. Genome Res. 2011;21:974–84.

Andrews S. FastQC: a quality control tool for high throughput sequence data. Available online at: http://www.bioinformatics.babraham.ac.uk/projects/fastqc. 2010.

Audano PA, Sulovari A, Graves-Lindsay TA, Cantsilieris S, Sorensen M, Welch AE, Dougherty ML, Nelson BJ, Shah A, Dutcher SK, Warren WC, Magrini V, McGrath SD, Li YI, Wilson RK, Eichler EE. Characterizing the Major Structural Variant Alleles of the Human Genome. Cell. 2019 Jan 24;176(3):663–675.e19. doi: 10.1016/j.cell.2018.12.019.

Balart-García P, Aristide L, Bradford TM, Beasley-Hall PG, Polak S, Cooper SJB, Fernández R. Parallel and convergent genomic changes underlie independent subterranean colonization across beetles. Nat Commun. 2023 Jun 29;14(1):3842. doi: 10.1038/s41467-023-39603-1.

Bilandžija H, Ma L, Parkhurst A, Jeffery WR. A potential benefit of albinism in Astyanax cavefish: downregulation of the oca2 gene increases tyrosine and catecholamine levels as an alternative to melanin synthesis. PLoS One. 2013 Nov 25;8(11):e80823. doi: 10.1371/journal.pone.0080823.

Boettger LM, Salem RM, Handsaker RE, Peloso GM, Kathiresan S, Hirschhorn JN, McCarroll SA. Recurring exon deletions in the HP (haptoglobin) gene contribute to lower blood cholesterol levels. Nat Genet. 2016 Apr;48(4):359–66. doi: 10.1038/ng.3510.

Bolger AM, Lohse M, Usadel B. Trimmomatic: a flexible trimmer for Illumina sequence data. Bioinformatics. 2014 Aug 1;30(15):2114–20. doi: 10.1093/bioinformatics/btu170.

Bradic M, Teotónio H, Borowsky RL. The population genomics of repeated evolution in the blind cavefish Astyanax mexicanus. Mol Biol Evol. 2013 Nov;30(11):2383–400. doi: 10.1093/molbev/mst136.

Bradic M, Beerli P, García-de León FJ, Esquivel-Bobadilla S, Borowsky RL. Gene flow and population structure in the Mexican blind cavefish complex (Astyanax mexicanus). BMC Evol Biol. 2012 Jan 23;12:9. doi: 10.1186/1471-2148-12-9.

Chain FJ, Feulner PG, Panchal M, Eizaguirre C, Samonte IE, Kalbe M, Lenz TL, Stoll M, Bornberg-Bauer E, Milinski M, Reusch TB. Extensive copy-number variation of young genes across stickleback populations. PLoS Genet. 2014 Dec 4;10(12):e1004830. doi: 10.1371/journal.pgen.1004830.

Dorant Y, Cayuela H, Wellband K, Laporte M, Rougemont Q, Mérot C, Normandeau E, Rochette R, Bernatchez L. Copy number variants outperform SNPs to reveal genotype-temperature association in a marine species. Mol Ecol. 2020 Dec;29(24):4765–4782. doi: 10.1111/mec.15565.

Duboué ER, Keene AC, Borowsky RL. Evolutionary convergence on sleep loss in cavefish populations. Curr Biol. 2011 Apr 26;21(8):671–6. doi: 10.1016/j.cub.2011.03.020.

Ebert P, Audano PA, Zhu Q, Rodriguez-Martin B, Porubsky D, Bonder MJ, Sulovari A, Ebler J, Zhou W, Serra Mari R, Yilmaz F, Zhao X, Hsieh P, Lee J, Kumar S, Lin J, Rausch T, Chen Y, Ren J, Santamarina M, Höps W, Ashraf H, Chuang NT, Yang X, Munson KM, Lewis AP, Fairley S, Tallon LJ, Clarke WE, Basile AO, Byrska-Bishop M, Corvelo A, Evani US, Lu TY, Chaisson MJP, Chen J, Li C, Brand H, Wenger AM, Ghareghani M, Harvey WT, Raeder B, Hasenfeld P, Regier AA, Abel HJ, Hall IM, Flicek P, Stegle O, Gerstein MB, Tubio JMC, Mu Z, Li YI, Shi X, Hastie AR, Ye K, Chong Z, Sanders AD, Zody MC, Talkowski ME, Mills RE, Devine SE, Lee C, Korbel JO, Marschall T, Eichler EE. Haplotype-resolved diverse human genomes and integrated analysis of structural variation. Science. 2021 Apr 2;372(6537):eabf7117. doi: 10.1126/science.abf7117.

Elipot Y, Hinaux H, Callebert J, Rétaux S. Evolutionary shift from fighting to foraging in blind cavefish through changes in the serotonin network. Curr Biol. 2013 Jan 7;23(1):1–10. doi: 10.1016/j.cub.2012.10.044.

Fumey J, Hinaux H, Noirot C, Thermes C, Rétaux S, Casane D. Evidence for late Pleistocene origin of Astyanax mexicanus cavefish. BMC Evol Biol. 2018 Apr 18;18(1):43. doi: 10.1186/s12862-018-1156-7.

Gamboa JL, Andrade FH. Mitochondrial content and distribution changes specific to mouse diaphragm after chronic normobaric hypoxia. Am J Physiol Regul Integr Comp Physiol. 2010 Mar;298(3):R575–83. doi: 10.1152/ajpregu.00320.2009.

Garg P, Martin-Trujillo A, Rodriguez OL, Gies SJ, Hadelia E, Jadhav B, Jain M, Paten B, Sharp AJ. Pervasive cis effects of variation in copy number of large tandem repeats on local DNA methylation and gene expression. Am J Hum Genet. 2021 May 6;108(5):809–824. doi: 10.1016/j.ajhg.2021.03.016.

Gene [Internet]. Bethesda (MD): National Library of Medicine (US), National Center for Biotechnology Information; [1988] –. Gene ID: 440243, GOLGA6L22 golgin A6 family like 22 [Homo sapiens (human)]; [cited 2023 07 20]. Available from: https://www.ncbi.nlm.nih.gov/gene/440243/#gene-expression

Gene [Internet]. Bethesda (MD): National Library of Medicine (US), National Center for Biotechnology Information; [1988] – Gene ID: 80833, APOL3 apolipoprotein L3 [Homo sapiens (human)]; [cited 2023 07 20]. Available from: https://www.ncbi.nlm.nih.gov/gene/80833

Gross JB. The complex origin of Astyanax cavefish. BMC Evol Biol. 2012 Jun 30;12:105. doi: 10.1186/1471-2148-12-105.

Gutsaeva DR, Carraway MS, Suliman HB, Demchenko IT, Shitara H, Yonekawa H, Piantadosi CA. Transient hypoxia stimulates mitochondrial biogenesis in brain subcortex by a neuronal nitric oxide synthase-dependent mechanism. J Neurosci. 2008 Feb 27;28(9):2015–24. doi: 10.1523/JNEUROSCI.5654-07.2008.

Herman A, Brandvain Y, Weagley J, Jeffery WR, Keene AC, Kono TJY, Bilandžija H, Borowsky R, Espinasa L, O’Quin K, Ornelas-García CP, Yoshizawa M, Carlson B, Maldonado E, Gross JB, Cartwright RA, Rohner N, Warren WC, McGaugh SE. The role of gene flow in rapid and repeated evolution of cave-related traits in Mexican tetra, Astyanax mexicanus. Mol Ecol. 2018 Nov;27(22):4397–4416. doi: 10.1111/mec.14877.

Hirase S, Ozaki H, Iwasaki W. Parallel selection on gene copy number variations through evolution of three-spined stickleback genomes. BMC Genomics. 2014 Aug 29;15(1):735. doi: 10.1186/1471-2164-15-735.

Huddleston J, Chaisson MJP, Steinberg KM, Warren W, Hoekzema K, Gordon D, Graves-Lindsay TA, Munson KM, Kronenberg ZN, Vives L, Peluso P, Boitano M, Chin CS, Korlach J, Wilson RK, Eichler EE. Discovery and genotyping of structural variation from long-read haploid genome sequence data. Genome Res. 2017 May;27(5):677–685. doi: 10.1101/gr.214007.116.

Ishikawa A, Yamanouchi S, Iwasaki W, Kitano J. Convergent copy number increase of genes associated with freshwater colonization in fishes. Philos Trans R Soc Lond B Biol Sci. 2022 Jul 18;377(1855):20200509. doi: 10.1098/rstb.2020.0509.

Iskow RC, Gokcumen O, Lee C. Exploring the role of copy number variants in human adaptation. Trends Genet. 2012 Jun;28(6):245–57. doi: 10.1016/j.tig.2012.03.002.

Jaggard JB, Stahl BA, Lloyd E, Prober DA, Duboue ER, Keene AC. Hypocretin underlies the evolution of sleep loss in the Mexican cavefish. Elife. 2018 Feb 6;7:e32637. doi: 10.7554/eLife.32637.

Jakt LM, Dubin A, Johansen SD. Intron size minimisation in teleosts. BMC Genomics. 2022 Sep 1;23(1):628. doi: 10.1186/s12864-022-08760-w.

Jang J, Terefe E, Kim K, Lee YH, Belay G, Tijjani A, Han JL, Hanotte O, Kim H. Population differentiated copy number variation of Bos taurus, Bos indicus and their African hybrids. BMC Genomics. 2021 Jul 12;22(1):531. doi: 10.1186/s12864-021-07808-7.

Jones FC, Grabherr MG, Chan YF, Russell P, Mauceli E, Johnson J, Swofford R, Pirun M, Zody MC, White S, Birney E, Searle S, Schmutz J, Grimwood J, Dickson MC, Myers RM, Miller CT, Summers BR, Knecht AK, Brady SD, Zhang H, Pollen AA, Howes T, Amemiya C; Broad Institute Genome Sequencing Platform & Whole Genome Assembly Team; Baldwin J, Bloom T, Jaffe DB, Nicol R, Wilkinson J, Lander ES, Di Palma F, Lindblad-Toh K, Kingsley DM. The genomic basis of adaptive evolution in threespine sticklebacks. Nature. 2012 Apr 4;484(7392):55–61. doi: 10.1038/nature10944.

Kondrashov FA, Rogozin IB, Wolf YI, Koonin EV. Selection in the evolution of gene duplications. Genome Biol. 2002;3(2):RESEARCH0008. doi: 10.1186/gb-2002-3-2-research0008.

Kondrashov FA. Gene duplication as a mechanism of genomic adaptation to a changing environment. Proc Biol Sci. 2012 Dec 22;279(1749):5048–57. doi: 10.1098/rspb.2012.1108.

Kosugi S, Momozawa Y, Liu X, Terao C, Kubo M, Kamatani Y. Comprehensive evaluation of structural variation detection algorithms for whole genome sequencing. Genome Biol. 2019 Jun 3;20(1):117. doi: 10.1186/s13059-019-1720-5.

Kowalko JE, Rohner N, Rompani SB, Peterson BK, Linden TA, Yoshizawa M, Kay EH, Weber J, Hoekstra HE, Jeffery WR, Borowsky R, Tabin CJ. Loss of schooling behavior in cavefish through sight-dependent and sight-independent mechanisms. Curr Biol. 2013 Oct 7;23(19):1874–83. doi: 10.1016/j.cub.2013.07.056.

Lai YT, Yeung CKL, Omland KE, Pang EL, Hao Y, Liao BY, Cao HF, Zhang BW, Yeh CF, Hung CM, Hung HY, Yang MY, Liang W, Hsu YC, Yao CT, Dong L, Lin K, Li SH. Standing genetic variation as the predominant source for adaptation of a songbird. Proc Natl Acad Sci U S A. 2019 Feb 5;116(6):2152–2157. doi: 10.1073/pnas.1813597116.

Langmead B, Salzberg SL. Fast gapped-read alignment with Bowtie 2. Nat Methods. 2012 Mar 4;9(4):357–9. doi: 10.1038/nmeth.1923.

Legendre L, Rode J, Germon I, Pavie M, Quiviger C, Policarpo M, Leclercq J, Père S, Fumey J, Hyacinthe C, Ornelas-García P, Espinasa L, Rétaux S, Casane D. Genetic identification and reiterated captures suggest that the Astyanax mexicanus El Pachón cavefish population is closed and declining. Zool Res. 2023 Jul 18;44(4):701–711. doi: 10.24272/j.issn.2095-8137.2022.481.

Li H, Handsaker B, Wysoker A, Fennell T, Ruan J, Homer N, Marth G, Abecasis G, Durbin R; 1000 Genome Project Data Processing Subgroup. The Sequence Alignment/Map format and SAMtools. Bioinformatics. 2009 Aug 15;25(16):2078–9. doi: 10.1093/bioinformatics/btp352.

Lowe CB, Sanchez-Luege N, Howes TR, Brady SD, Daugherty RR, Jones FC, Bell MA, Kingsley DM. Detecting differential copy number variation between groups of samples. Genome Res. 2018 Feb;28(2):256–265. doi: 10.1101/gr.206938.116.

Maron LG, Guimarães CT, Kirst M, Albert PS, Birchler JA, Bradbury PJ, Buckler ES, Coluccio AE, Danilova TV, Kudrna D, Magalhaes JV, Piñeros MA, Schatz MC, Wing RA, Kochian LV. Aluminum tolerance in maize is associated with higher MATE1 gene copy number. Proc Natl Acad Sci U S A. 2013 Mar 26;110(13):5241–6. doi: 10.1073/pnas.1220766110.

Martin M. Cutadapt removes adapter sequences from high-throughput sequencing reads. EMBnet. J. 2011;17:10–12. doi: 10.14806/ej.17.1.200.

Meng G, Bao Q, Ma X, Chu M, Huang C, Guo X, Liang C, Yan P. Analysis of Copy Number Variation in the Whole Genome of Normal-Haired and Long-Haired Tianzhu White Yaks. Genes (Basel). 2022 Dec 18;13(12):2405. doi: 10.3390/genes13122405.

Menuet A, Alunni A, Joly JS, Jeffery WR, Rétaux S. Expanded expression of Sonic Hedgehog in Astyanax cavefish: multiple consequences on forebrain development and evolution. Development. 2007 Mar;134(5):845–55. doi: 10.1242/dev.02780.

Mölder F, Jablonski KP, Letcher B, Hall MB, Tomkins-Tinch CH, Sochat V, Forster J, Lee S, Twardziok SO, Kanitz A, Wilm A, Holtgrewe M, Rahmann S, Nahnsen S, Köster J. Sustainable data analysis with Snakemake. F1000Res. 2021 Jan 18;10:33. doi: 10.12688/f1000research.29032.2.

Moran D, Softley R, Warrant EJ. The energetic cost of vision and the evolution of eyeless Mexican cavefish. Science Advances. 2015;1:e1500363. doi: 10.1126/sciadv.1500363.

Niederreither K, Vermot J, Fraulob V, Chambon P, Dolle P. Retinaldehyde dehydrogenase 2 (RALDH2)-independent patterns of retinoic acid synthesis in the mouse embryo. Proc Natl Acad Sci U S A. 2002 Dec 10;99(25):16111–6. doi: 10.1073/pnas.252626599.

Open2C, Nezar Abdennur, Geoffrey Fudenberg, Ilya Flyamer, Aleksandra A. Galitsyna, Anton Goloborodko, Maxim Imakaev, Sergey V. Venev: Bioframe: Operations on Genomic Intervals in Pandas Dataframes. bioRxiv. Preprint. 2022.02.16.480748; doi: 10.1101/2022.02.16.480748

Peuß R, Box AC, Chen S, Wang Y, Tsuchiya D, Persons JL, Kenzior A, Maldonado E, Krishnan J, Scharsack JP, Slaughter BD, Rohner N. Adaptation to low parasite abundance affects immune investment and immunopathological responses of cavefish. Nat Ecol Evol. 2020 Oct;4(10):1416–1430. doi: 10.1038/s41559-020-1234-2.

Pezer Ž, Harr B, Teschke M, Babiker H, Tautz D. Divergence patterns of genic copy number variation in natural populations of the house mouse (Mus musculus domesticus) reveal three conserved genes with major population-specific expansions. Genome Res. 2015 Aug;25(8):1114–24. doi: 10.1101/gr.187187.114.

Riddle MR, Aspiras AC, Gaudenz K, Peuß R, Sung JY, Martineau B, Peavey M, Box AC, Tabin JA, McGaugh S, Borowsky R, Tabin CJ, Rohner N. Insulin resistance in cavefish as an adaptation to a nutrient-limited environment. Nature. 2018 Mar 29;555(7698):647-651. doi: 10.1038/nature26136.

Rinker DC, Specian NK, Zhao S, Gibbons JG. Polar bear evolution is marked by rapid changes in gene copy number in response to dietary shift. Proc Natl Acad Sci U S A. 2019 Jul 2;116(27):13446–13451. doi: 10.1073/pnas.1901093116.

Rundle HD, Nagel L, Wenrick Boughman J, Schluter D. Natural selection and parallel speciation in sympatric sticklebacks. Science. 2000 Jan 14;287(5451):306-8. doi: 10.1126/science.287.5451.306.

Saitou M, Masuda N, Gokcumen O. Similarity-Based Analysis of Allele Frequency Distribution among Multiple Populations Identifies Adaptive Genomic Structural Variants. Mol Biol Evol. 2022 Mar 2;39(3):msab313. doi: 10.1093/molbev/msab313.

Sarkar A, Sim C, Hong YS, Hogan JR, Fraser MJ, Robertson HM, Collins FH. Molecular evolutionary analysis of the widespread piggyBac transposon family and related “domesticated” sequences. Mol Genet Genomics. 2003 Nov;270(2):173–80. doi: 10.1007/s00438-003-0909-0.

Shebanits K, Günther T, Johansson ACV, Maqbool K, Feuk L, Jakobsson M, Larhammar D. Copy number determination of the gene for the human pancreatic polypeptide receptor NPY4R using read depth analysis and droplet digital PCR. BMC Biotechnol. 2019 Jun 4;19(1):31. doi: 10.1186/s12896-019-0523-9.

Sherman BT, Hao M, Qiu J, Jiao X, Baseler MW, Lane HC, Imamichi T, Chang W. DAVID: a web server for functional enrichment analysis and functional annotation of gene lists (2021 update). Nucleic Acids Res. 2022 Mar 23;50(W1):W216–21. doi: 10.1093/nar/gkac194.

Singh G, Davenport AP. Neuropeptide B and W: neurotransmitters in an emerging G-protein-coupled receptor system. Br J Pharmacol. 2006 Aug;148(8):1033–41. doi: 10.1038/sj.bjp.0706825.

Solé M, Ablondi M, Binzer-Panchal A, Velie BD, Hollfelder N, Buys N, Ducro BJ, François L, Janssens S, Schurink A, Viklund Å, Eriksson S, Isaksson A, Kultima H, Mikko S, Lindgren G. Inter- and intra-breed genome-wide copy number diversity in a large cohort of European equine breeds. BMC Genomics. 2019 Oct 22;20(1):759. doi: 10.1186/s12864-019-6141-z.

Suvakov M, Panda A, Diesh C, Holmes I, Abyzov A. CNVpytor: a tool for copy number variation detection and analysis from read depth and allele imbalance in whole-genome sequencing. Gigascience. 2021 Nov 18;10(11):giab074. doi: 10.1093/gigascience/giab074.

tan Nguyen H, Merriman TR, Black MA. CNVrd, a read-depth algorithm for assigning copy-number at the FCGR locus: population-specific tagging of copy number variation at FCGR3B. PLoS One. 2013 Apr 30;8(4):e63219. doi: 10.1371/journal.pone.0063219.

Uhlén M, Fagerberg L, Hallström BM, Lindskog C, Oksvold P, Mardinoglu A, Sivertsson Å, Kampf C, Sjöstedt E, Asplund A, Olsson I, Edlund K, Lundberg E, Navani S, Szigyarto CA, Odeberg J, Djureinovic D, Takanen JO, Hober S, Alm T, Edqvist PH, Berling H, Tegel H, Mulder J, Rockberg J, Nilsson P, Schwenk JM, Hamsten M, von Feilitzen K, Forsberg M, Persson L, Johansson F, Zwahlen M, von Heijne G, Nielsen J, Pontén F. Proteomics. Tissue-based map of the human proteome. Science. 2015 Jan 23;347(6220):1260419. doi: 10.1126/science.1260419.

van der Weele CM, Jeffery WR. Cavefish cope with environmental hypoxia by developing more erythrocytes and overexpression of hypoxia-inducible genes. Elife. 2022 Jan 5;11:e69109. doi: 10.7554/eLife.69109.

Vickrey AI, Bruders R, Kronenberg Z, Mackey E, Bohlender RJ, Maclary ET, Maynez R, Osborne EJ, Johnson KP, Huff CD, Yandell M, Shapiro MD. Introgression of regulatory alleles and a missense coding mutation drive plumage pattern diversity in the rock pigeon. Elife. 2018 Jul 17;7:e34803. doi: 10.7554/eLife.34803.

Warren WC, Boggs TE, Borowsky R, Carlson BM, Ferrufino E, Gross JB, Hillier L, Hu Z, Keene AC, Kenzior A, Kowalko JE, Tomlinson C, Kremitzki M, Lemieux ME, Graves-Lindsay T, McGaugh SE, Miller JT, Mommersteeg MTM, Moran RL, Peuß R, Rice ES, Riddle MR, Sifuentes-Romero I, Stanhope BA, Tabin CJ, Thakur S, Yamamoto Y, Rohner N. A chromosome-level genome of Astyanax mexicanus surface fish for comparing population-specific genetic differences contributing to trait evolution. Nat Commun. 2021 Mar 4;12(1):1447. doi: 10.1038/s41467-021-21733-z.

Warren WC, Carroll RA, Haggerty L, Keene AC, McGaugh SE, Ogeh D, Rice ES, Roback E, Rohner N, Martin F, Maggs X. Astyanax mexicanus surface and cavefish chromosome-scale assemblies for trait variation discovery. bioRxiv 2023.11.16.567450; doi: 10.1101/2023.11.16.567450

Wozniak D, Quinnell T. Unmet needs of patients with narcolepsy: perspectives on emerging treatment options. Nat Sci Sleep. 2015;7:51–61. 10.2147/NSS.S56077

Xiong S, Krishnan J, Peuß R, Rohner N. Early adipogenesis contributes to excess fat accumulation in cave populations of Astyanax mexicanus. Dev Biol. 2018 Sep 15;441(2):297–304. doi: 10.1016/j.ydbio.2018.06.003.

Xu L, Hou Y, Bickhart DM, Zhou Y, Hay el HA, Song J, Sonstegard TS, Van Tassell CP, Liu GE. Population-genetic properties of differentiated copy number variations in cattle. Sci Rep. 2016 Mar 23;6:23161. doi: 10.1038/srep23161.

Yang L, Han J, Deng T, Li F, Han X, Xia H, Quan F, Hua G, Yang L, Zhou Y. Comparative analyses of copy number variations between swamp buffaloes and river buffaloes. Anim Genet. 2023 Apr;54(2):199–206. doi: 10.1111/age.13288.

Yi G, Qu L, Liu J, Yan Y, Xu G, Yang N. Genome-wide patterns of copy number variation in the diversified chicken genomes using next-generation sequencing. BMC Genomics. 2014 Nov 7;15(1):962. doi: 10.1186/1471-2164-15-962.

Yoshizawa M, Goricki S, Soares D, Jeffery WR. Evolution of a behavioral shift mediated by superficial neuromasts helps cavefish find food in darkness. Curr Biol. 2010 Sep 28;20(18):1631–6. doi: 10.1016/j.cub.2010.07.017.

Yuste-Lisbona FJ, Fernández-Lozano A, Pineda B, Bretones S, Ortíz-Atienza A, García-Sogo B, Müller NA, Angosto T, Capel J, Moreno V, Jiménez-Gómez JM, Lozano R. ENO regulates tomato fruit size through the floral meristem development network. Proc Natl Acad Sci U S A. 2020 Apr 7;117(14):8187–8195. doi: 10.1073/pnas.1913688117.

Zhang F, Gu W, Hurles ME, Lupski JR. Copy number variation in human health, disease, and evolution. Annu Rev Genomics Hum Genet. 2009;10:451–81. doi: 10.1146/annurev.genom.9.081307.164217.

Zhu C, Li M, Qin S, Zhao F, Fang S. Detection of copy number variation and selection signatures on the X chromosome in Chinese indigenous sheep with different types of tail. Asian-Australas J Anim Sci. 2020 Sep;33(9):1378–1386. doi: 10.5713/ajas.18.0661.

Zong SB, Li YL, Liu JX. Genomic Architecture of Rapid Parallel Adaptation to Fresh Water in a Wild Fish. Mol Biol Evol. 2021 Apr 13;38(4):1317–1329. doi: 10.1093/molbev/msaa290.

