## Supplementary Material for "The prevalence of copy number increase at multiallelic CNVs associated with cave colonization in Mexican tetra (*Astyanax mexicanus*)"

**Table S1.** Sequencing sample information (from Herman et al. 2018)

| SRR ID | Sample | Region | Habitat | Lineage |
|---|---|---|---|---|
| SRR1575288 | Molino2a | Molino | cave | new |
| SRR1575289 | Molino7a | Molino | cave | new |
| SRR1575290 | Molino9b | Molino | cave | new |
| SRR1575291 | Molino10b | Molino | cave | new |
| SRR1575292 | Molino11a | Molino | cave | new |
| SRR1575293 | Molino12a | Molino | cave | new |
| SRR1575294 | Molino13b | Molino | cave | new |
| SRR1575295 | Molino14a | Molino | cave | new |
| SRR1575296 | Molino15b | Molino | cave | new |
| SRR1575279 | Pach3 | Pachon | cave | old |
| SRR1575280 | Pach7 | Pachon | cave | old |
| SRR1575281 | Pach8 | Pachon | cave | old |
| SRR1575282 | Pach9 | Pachon | cave | old |
| SRR1575283 | Pach11 | Pachon | cave | old |
| SRR1575284 | Pach12 | Pachon | cave | old |
| SRR1575285 | Pach14 | Pachon | cave | old |
| SRR1575286 | Pach15 | Pachon | cave | old |
| SRR1575287 | Pach17 | Pachon | cave | old |
| SRR1927212 | Tinaja6 | Tinaja | cave | old |
| SRR1927214 | Tinaja12 | Tinaja | cave | old |
| SRR1927215 | TinajaB | Tinaja | cave | old |
| SRR1927218 | Tinaja2 | Tinaja | cave | old |
| SRR1927221 | TinajaC | Tinaja | cave | old |
| SRR1927224 | Tinaja3 | Tinaja | cave | old |
| SRR1927228 | TinajaD | Tinaja | cave | old |
| SRR1927232 | Tinaja5 | Tinaja | cave | old |
| SRR1927233 | TinajaE | Tinaja | cave | old |
| SRR1927184 | Tinaja11 | Tinaja | cave | old |
| SRR1575297 | Rascon02 | Rascon | surface | old |
| SRR1575298 | Rascon04 | Rascon | surface | old |
| SRR1927234 | Rascon13 | Rascon | surface | old |
| SRR1927235 | Rascon15 | Rascon | surface | old |
| SRR1927236 | Rascon8 | Rascon | surface | old |
| SRR1927237 | Rascon6 | Rascon | surface | old |
| SRR1575270 | Choy01 | Río Choy | surface | new |
| SRR1575271 | Choy05 | Río Choy | surface | new |
| SRR1575272 | Choy06 | Río Choy | surface | new |
| SRR1575273 | Choy09 | Río Choy | surface | new |
| SRR1575274 | Choy10 | Río Choy | surface | new |
| SRR1575275 | Choy11 | Río Choy | surface | new |
| SRR1575276 | Choy12 | Río Choy | surface | new |
| SRR1575277 | Choy13 | Río Choy | surface | new |
| SRR1575278 | Choy14 | Río Choy | surface | new |

**Table S2.** Per-base coverage and the number of detected CNVs

| Sample | Mean Base Coverage | CNV count | Deletions | Duplications |
|---|---|---|---|---|
| Molino2a | 9.8 | 5,923 | 5,481 (92.5 %) | 442 (7.5 %) |
| Molino7a | 9.4 | 5,216 | 4,885 (93.7 %) | 331 (6.3 %) |
| Molino9b | 9.0 | 5,073 | 4,749 (93.6 %) | 324 (6.4 %) |
| Molino10b | 10.1 | 5,942 | 5,559 (93.6 %) | 383 (6.4 %) |
| Molino11a | 8.1 | 4,721 | 4,348 (92.1 %) | 373 (7.9 %) |
| Molino12a | 9.0 | 4,977 | 4,619 (92.8 %) | 358 (7.2 %) |
| Molino13b | 11.8 | 7,353 | 6,873 (93.5 %) | 480 (6.5 %) |
| Molino14a | 7.5 | 4,291 | 3,935 (91.7 %) | 356 (8.3 %) |
| Molino15b | 7.6 | 4,462 | 4,094 (91.8 %) | 368 (8.2 %) |
| Pach3 | 11.0 | 7,575 | 7,253 (95.7 %) | 322 (4.3 %) |
| Pach7 | 8.3 | 4,799 | 4,561 (95.0 %) | 238 (5.0 %) |
| Pach8 | 9.0 | 5,067 | 4,829 (95.3 %) | 238 (4.7 %) |
| Pach9 | 8.4 | 5,306 | 5,072 (95.6 %) | 234 (4.4 %) |
| Pach11 | 7.7 | 4,814 | 4,571 (95.0 %) | 243 (5.0 %) |
| Pach12 | 9.3 | 5,602 | 5,361 (95.7 %) | 241 (4.3 %) |
| Pach14 | 7.7 | 4,926 | 4,706 (95.5 %) | 220 (4.5 %) |
| Pach15 | 9.4 | 5,721 | 5,492 (96.0 %) | 229 (4.0 %) |
| Pach17 | 7.7 | 4,743 | 4,517 (95.2 %) | 226 (4.8 %) |
| PachonRef | 17.7 | 3,648 | 3,263 (89.4 %) | 385 (10.6 %) |
| Tinaja6 | 8.9 | 4,558 | 4,345 (95.3 %) | 213 (4.7 %) |
| Tinaja12 | 11.2 | 8,414 | 8,077 (96.0 %) | 337 (4.0 %) |
| TinajaB | 10.7 | 8,024 | 7,692 (95.9 %) | 332 (4.1 %) |
| Tinaja2 | 8.0 | 5,438 | 5,159 (94.9 %) | 279 (5.1 %) |
| TinajaC | 10.0 | 6,847 | 6,570 (96.0 %) | 277 (4.0 %) |
| Tinaja3 | 8.9 | 5,657 | 5,380 (95.1 %) | 277 (4.9 %) |
| TinajaD | 8.0 | 5,391 | 5,157 (95.7 %) | 234 (4.3 %) |
| Tinaja5 | 6.6 | 4,527 | 4,277 (94.5 %) | 250 (5.5 %) |
| TinajaE | 9.6 | 6,548 | 6,316 (96.5 %) | 232 (3.5 %) |
| Tinaja11 | 10.1 | 6,861 | 6,601 (96.2 %) | 260 (3.8 %) |
| Rascon02 | 12.2 | 12,262 | 11,981 (97.7 %) | 281 (2.3 %) |
| Rascon04 | 11.5 | 9,982 | 9,745 (97.6 %) | 237 (2.4 %) |
| Rascon13 | 9.3 | 6,735 | 6,578 (97.7 %) | 157 (2.3 %) |
| Rascon15 | 9.6 | 7,586 | 7,405 (97.6 %) | 181 (2.4 %) |
| Rascon8 | 8.0 | 6,221 | 6,068 (97.5 %) | 153 (2.5 %) |
| Rascon6 | 9.1 | 6,167 | 6,019 (97.6 %) | 148 (2.4 %) |
| Choy01 | 8.8 | 3,143 | 2,850 (90.7 %) | 293 (9.3 %) |
| Choy05 | 9.7 | 3,390 | 3,099 (91.4 %) | 291 (8.6 %) |
| Choy06 | 8.2 | 3,509 | 3,215 (91.6 %) | 294 (8.4 %) |
| Choy09 | 8.9 | 3,241 | 2,932 (90.5 %) | 309 (9.5 %) |
| Choy10 | 8.6 | 3,001 | 2,723 (90.7 %) | 278 (9.3 %) |
| Choy11 | 9.1 | 3,195 | 2,883 (90.2 %) | 312 (9.8 %) |
| Choy12 | 9.2 | 3,203 | 2,921 (91.2 %) | 282 (8.8 %) |
| Choy13 | 9.8 | 3,824 | 3,468 (90.7 %) | 356 (9.3 %) |
| Choy14 | 8.6 | 3,055 | 2,786 (91.2 %) | 269 (8.8 %) |

**Figure S1.**

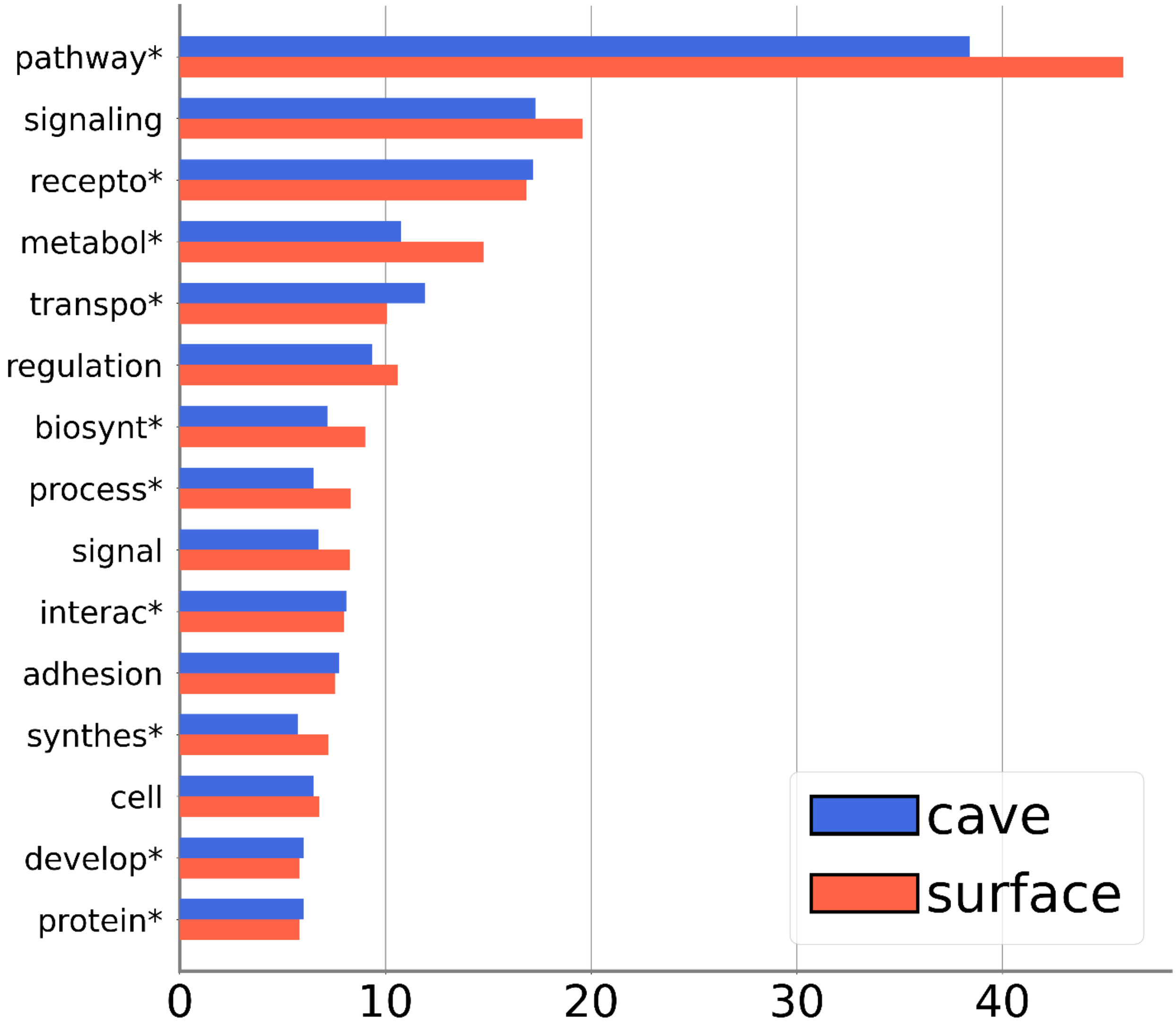


**Figure S1**. Most frequently occurring terms associated with genes that overlap CNVRs, analyzed separately for CNVRs found exclusively in cave (blue) or surface (red) ecotypes. Terms are derived from biological process and pathway annotations in DAVID tool. Proportions are shown as percentage of the number of genes associated with a term relative to the total number of genes that have assigned annotations in DAVID.

**Figure S2.**

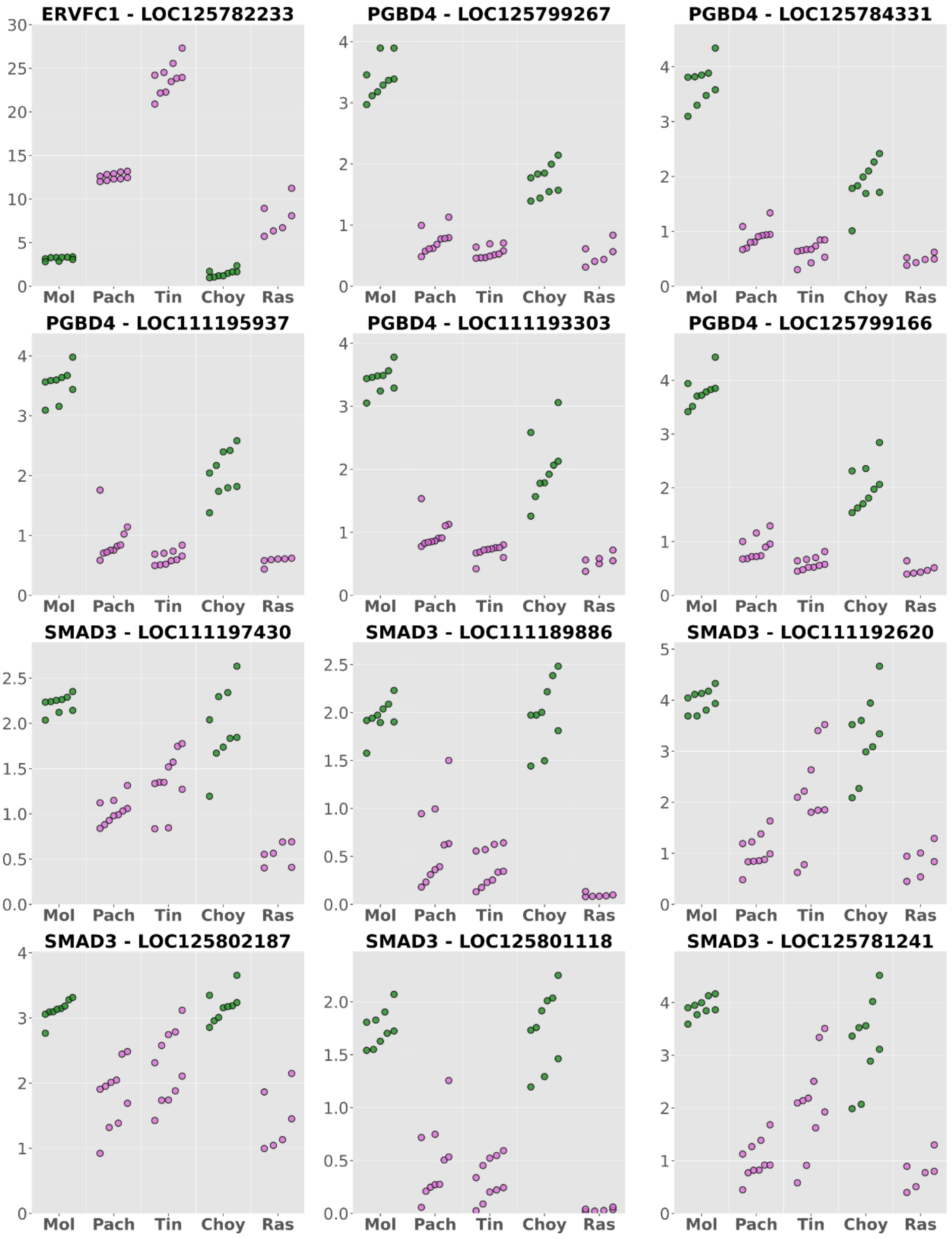


**Figure S2.** Copy numbers of genes ERVFC1, PGBD4 and SMAD3 annotated at genomic locations with significant differences between old (purple) and new (green) lineage individuals. Gene symbol and Gene ID are indicated on top of each plot. Mol - Molino; Pach - Pachon; Tin - Tinaja; Choy - Río Choy; Ras - Rascon.

**Figure S3.**

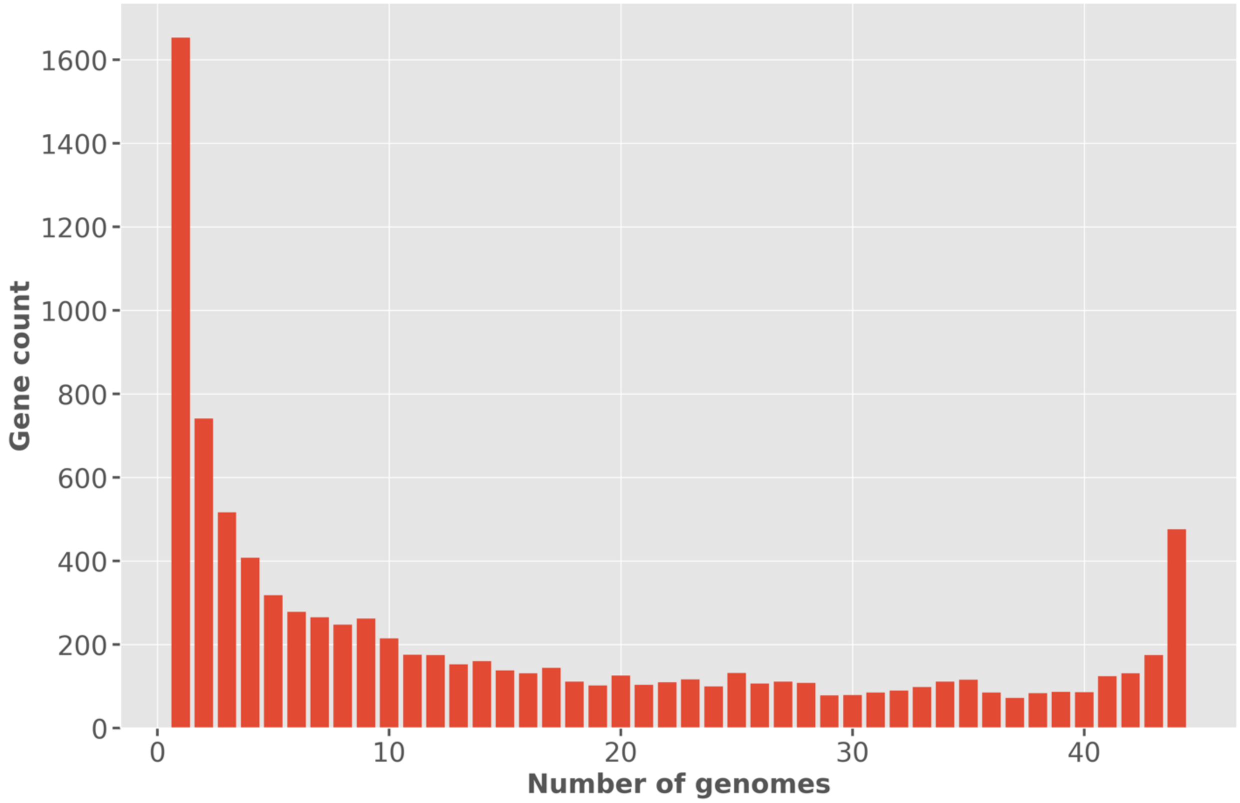


**Figure S3.** Number of protein-coding genes (Y-axis) versus the number of individuals (genomes) in which they intersect one or more CNVs.

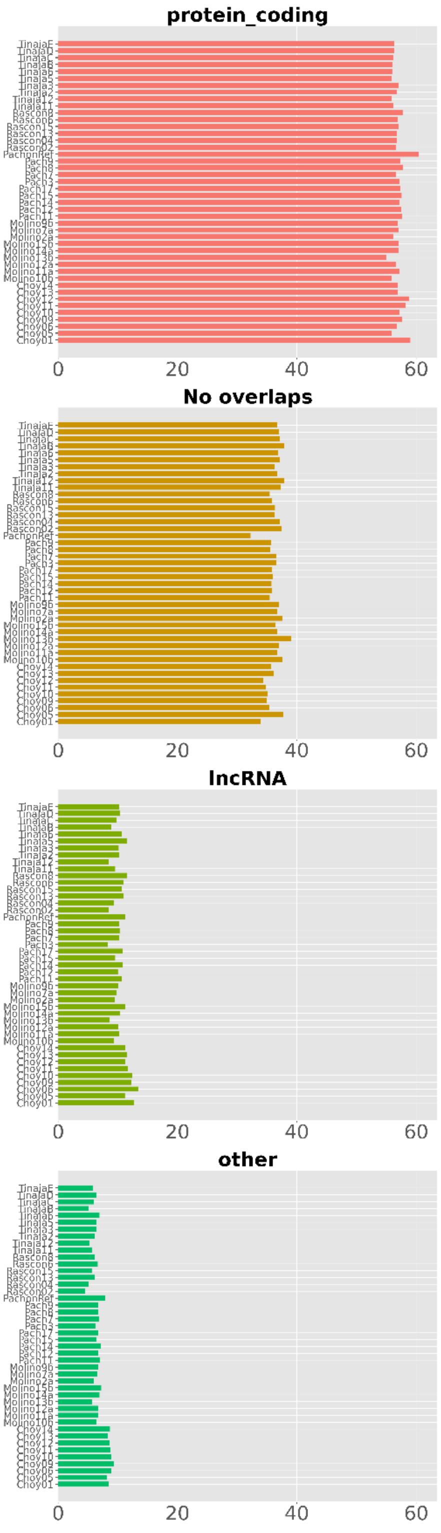


**Figure S4.** Proportion of CNVs affecting different gene categories per individual genome.

**Figure S5.**

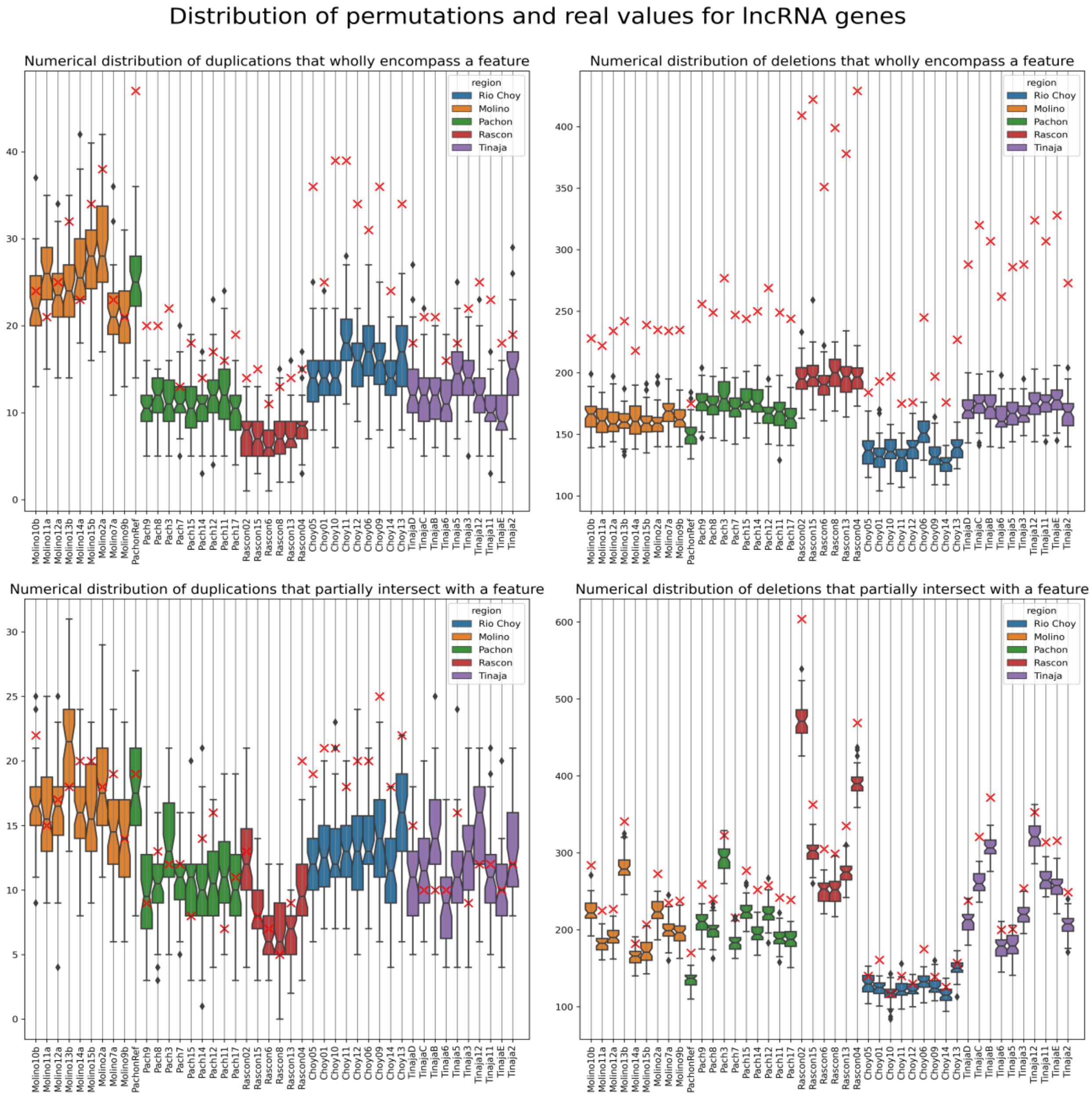
Distribution of permutations and real values for lncRNA genes
Numerical distribution of duplications that wholly encompass a feature
Numerical distribution of deletions that wholly encompass a feature
Numerical distribution of duplications that partially intersect with a feature
Numerical distribution of deletions that partially intersect with a feature
region
Rio Choy
Molino
Pachon
Rascon
Tinaja

**Figure S5.** Continued

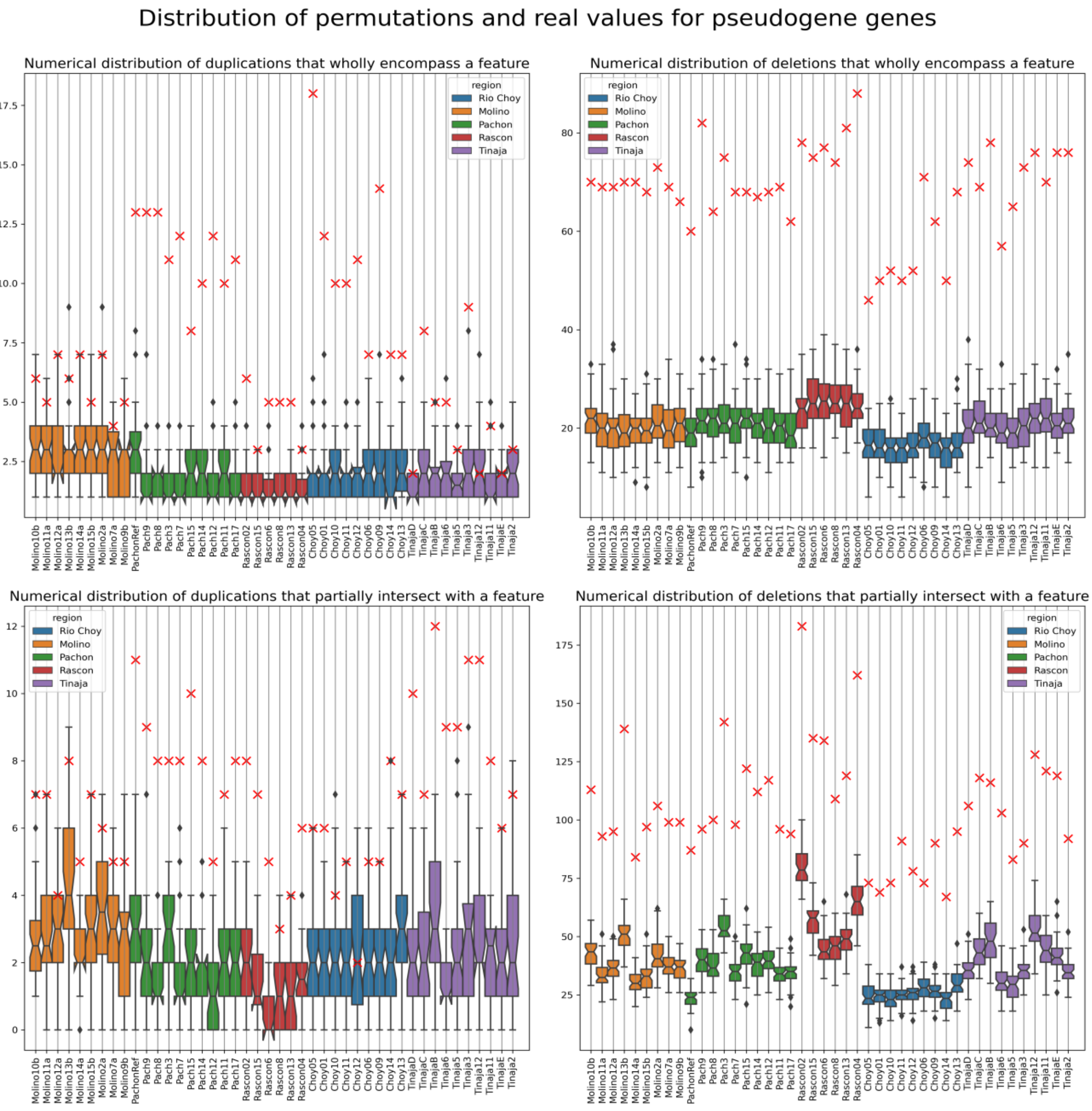

**Figure S5.** Continued

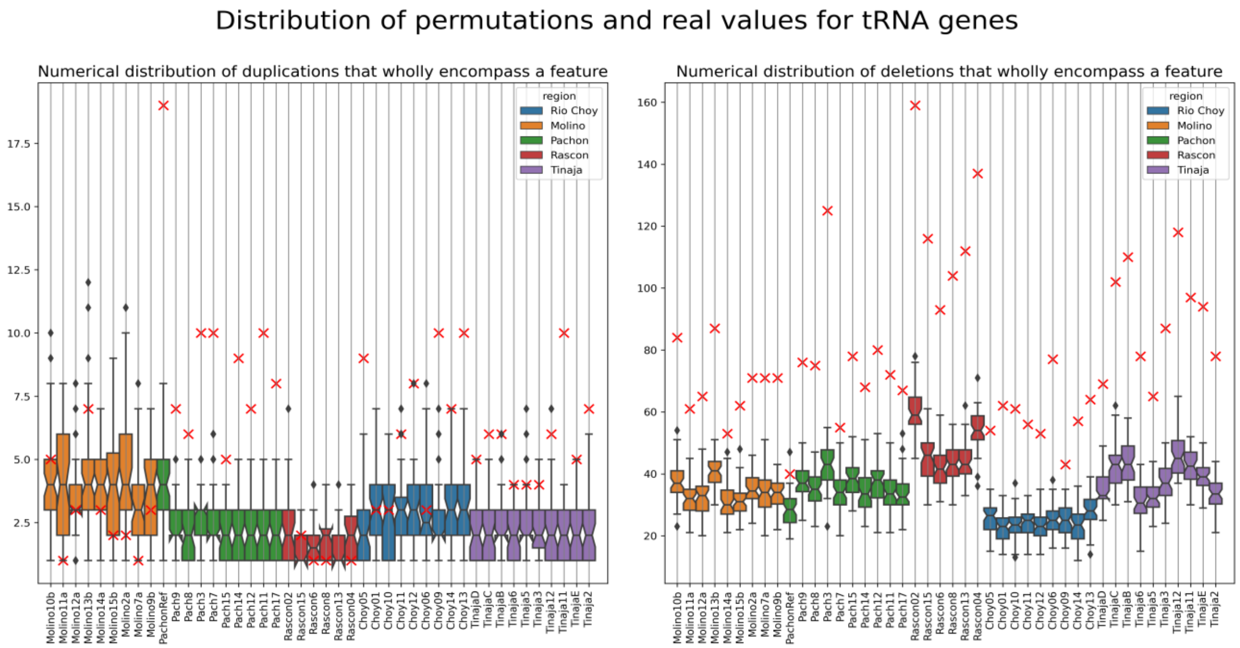


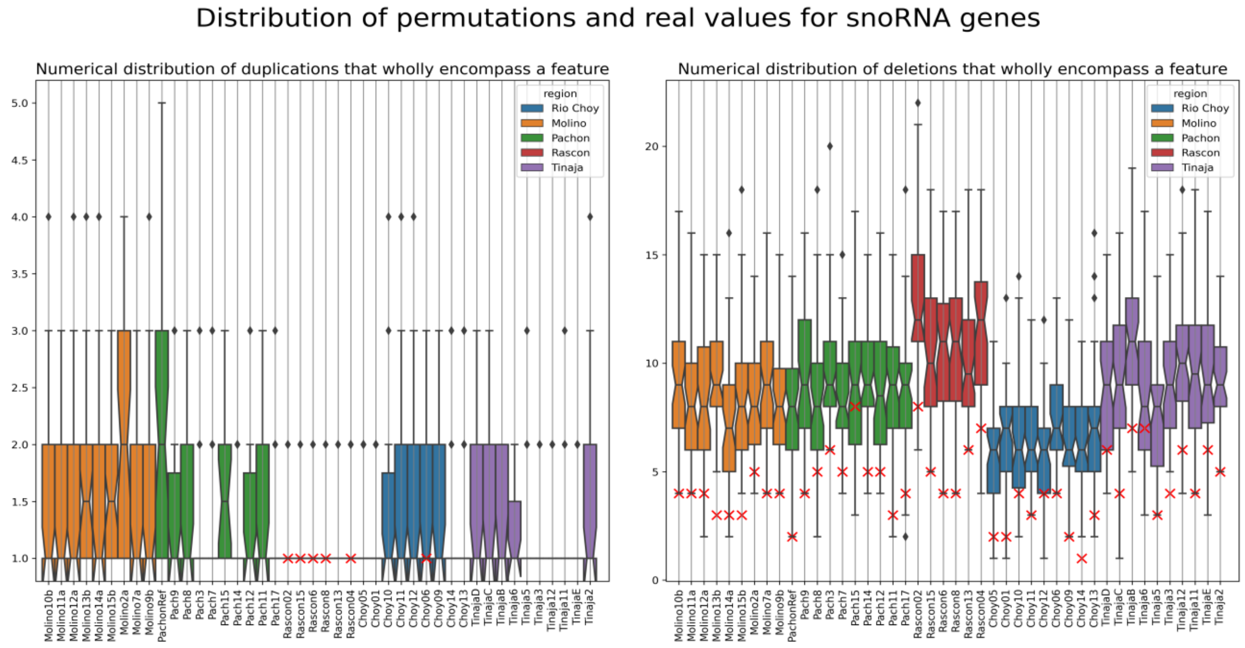

**Figure S5.** Continued

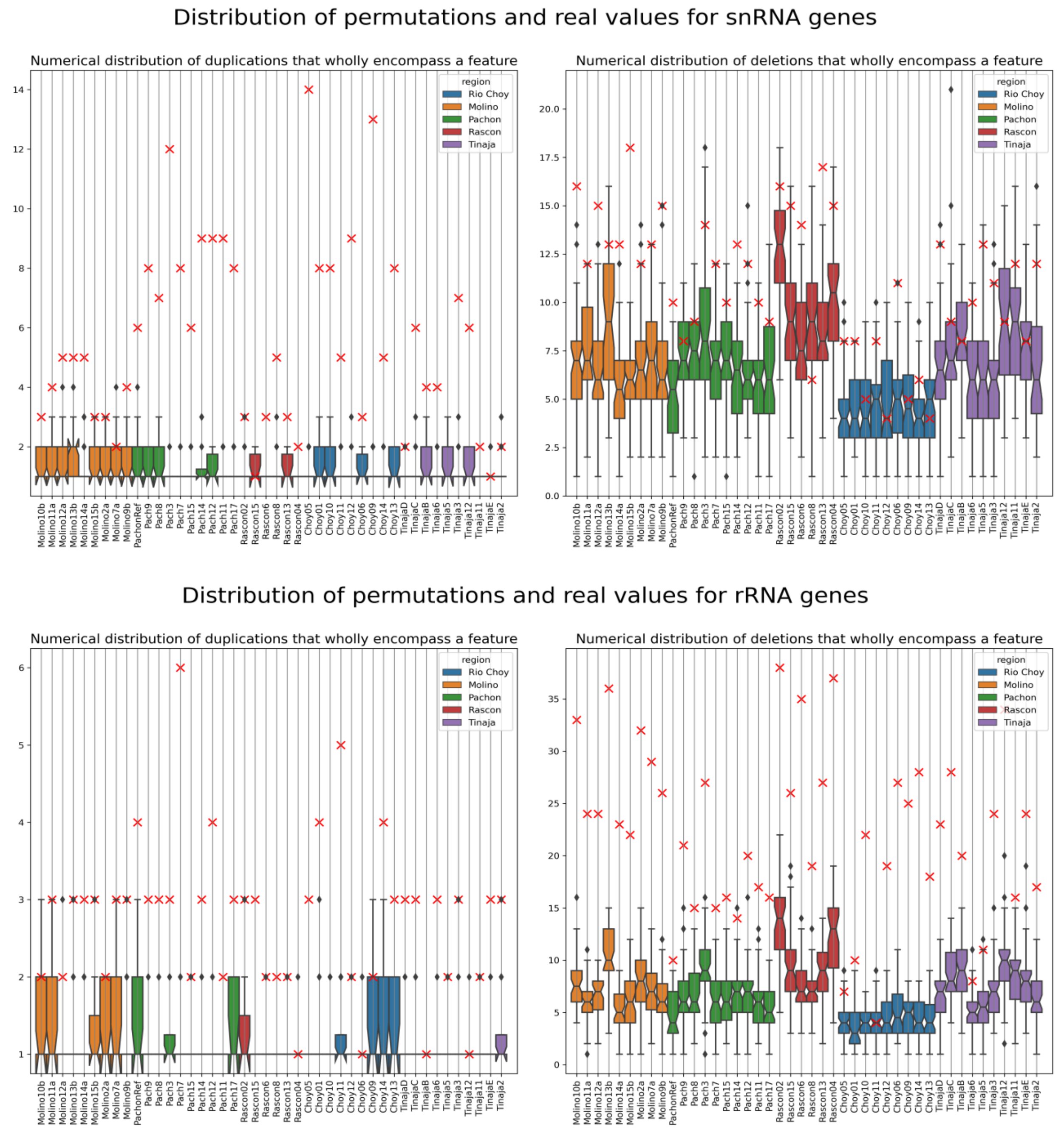


**Figure S5.** Number of CNVs that overlap annotated gene category (indicated in the plot title) detected in the true data set (red crosses), and in permuted data (distribution of expected values presented as boxplot), based on 50 permutations per sample.

**Figure S6.**

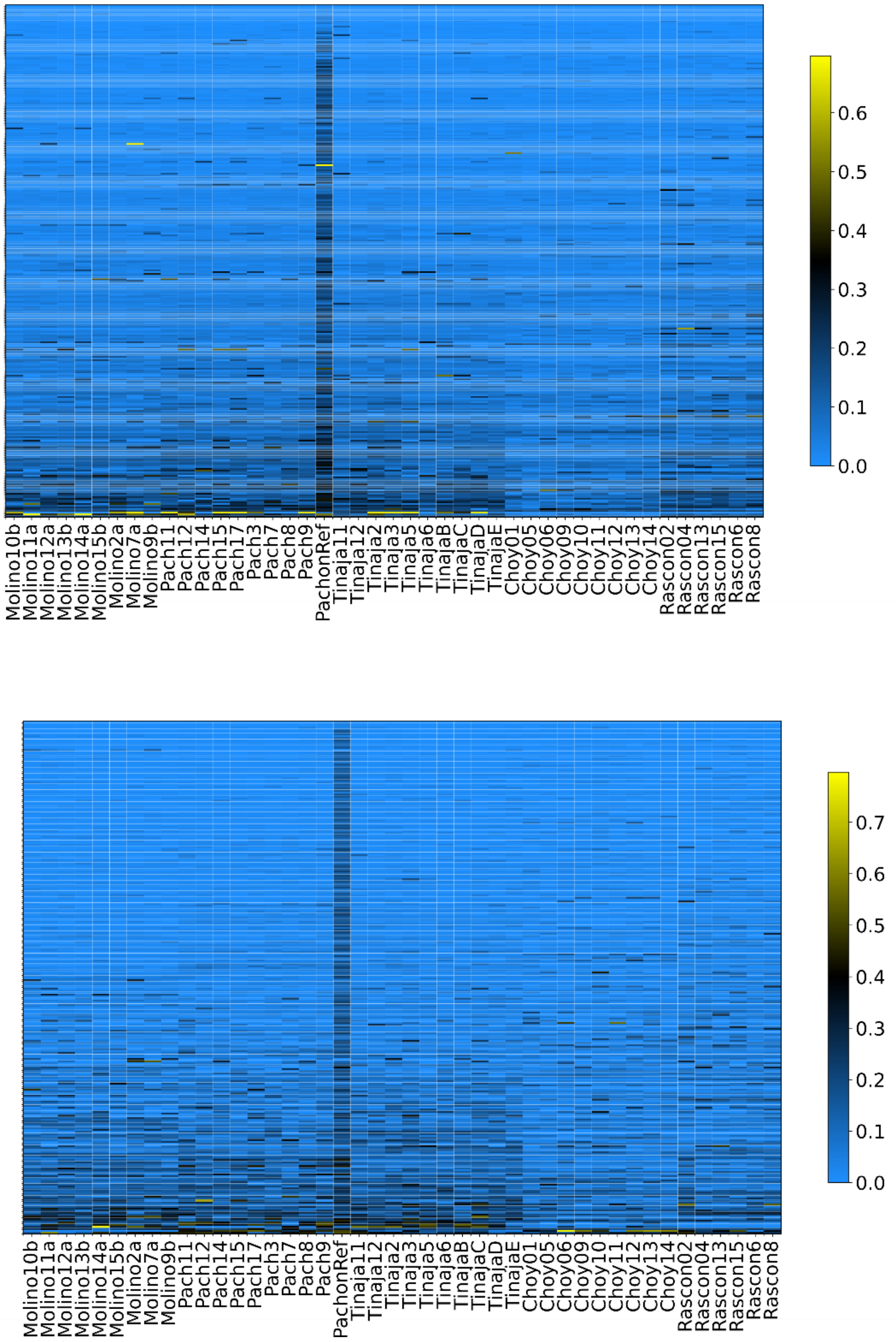


**Figure S6.** The fraction of reads with MAPQ=0 across each of the 292 CNVRs (upper panel) and 102 CNV genes (lower panel) that are divergent in CN between cave and surface fish. The fraction is calculated as the ratio between the number of reads with zero MAPQ and all reads mapped to the indicated region. The brightest yellow indicates maximal fraction within the dataset and the brightest blue indicates that there are no mapped reads with MAP=0.
